# High-throughput investigation of genetic design constraints in domesticated Influenza A Virus for transient gene delivery

**DOI:** 10.1101/2024.02.14.580300

**Authors:** Adam Sychla, Christopher S Stach, Shanley N Roach, Amanda N Hayward, Ryan A Langlois, Michael J Smanski

## Abstract

Replication-incompetent single cycle infectious Influenza A Virus (sciIAV) has demonstrated utility as a research and vaccination platform. Protein-based therapeutics are increasingly attractive due to their high selectivity and potent efficacy but still suffer from low bioavailability and high manufacturing cost. Transient RNA-mediated delivery is a safe alternative that allows for expression of protein-based therapeutics within the target cells or tissues but is limited by delivery efficiency. Here, we develop recombinant sciIAV as a platform for transient gene delivery *in vivo* and *in vitro* for therapeutic, research, and manufacturing applications (*in vivo* antimicrobial production, cell culture contamination clearance, and production of antiviral proteins *in vitro*). While adapting the system to deliver new protein cargo we discovered expression differences presumably resulting from genetic context effects. We applied a high-throughput screen to map these within the 3^*′*^-untranslated and coding regions of the hemagglutinin-encoding segment 4. This screen revealed permissible mutations in the 3^*′*^-UTR and depletion of RNA level motifs in the N-terminal coding region.

## Introduction

Throughout history, humans have refined and reshaped nature to create tools and technologies with desirable properties. Here, we explore and report several applications of domesticated Influenza A Virus (IAV).

IAV is a 174 MDa segmented, negative sense, single-stranded RNA virus that parasitizes human and other animal cells to replicate and spread (Figure 1a). In doing so, it causes between 9 and 41 million influenza cases per year in the US (1, 2). Structurally, IAV particles are composed of a lipid capsule co-opted from host cells. This capsule is decorated by 400-500 hemagglutinin (HA) and neuraminidase (NA) glycoproteins that are used for viral budding from and entry to host cells (1, 3). The weak 3 mM affinity of HA to sialic acid means that multivalent binding is likely required for proper attachment to cells (4). Due to this property, the virus evolved such that the glycoproteins comprise 50% of each particle’s mass (89 MDa) (1). As such, IAV particles are exquisitely optimized to produce large amounts of HA upon entry into host cells.

**Fig. 1.**
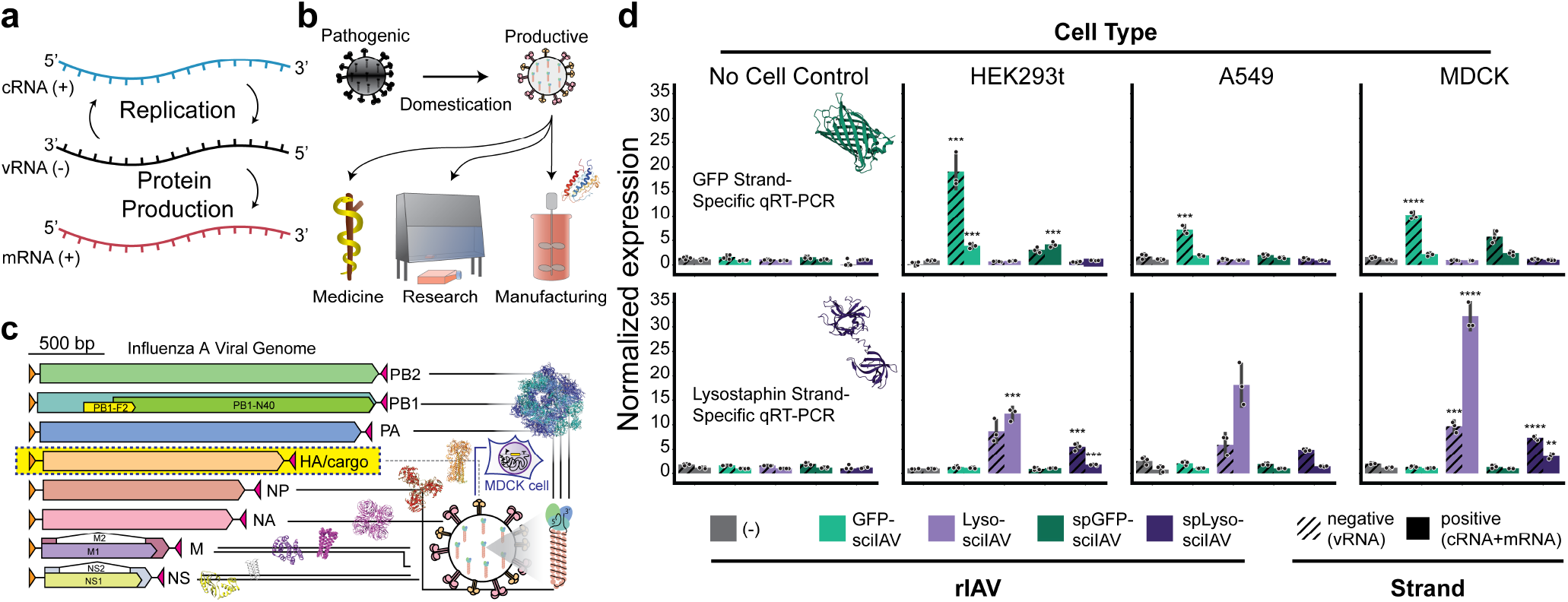
Domestication of IAV and confirmation of RNA expression. **(a)** RNA life cycle of IAV going through three RNA species for replication and protein production **(b)** Domestication of IAV removes pathogenicity and enables application in medicine, research, and manufacturing. **(c)** Genomic structure of IAV and domestication strategy. Each of eight genomic segments encodes one or more proteins, flanked by unique UTRs. The HA coding region can be replaced by cargo and HA protein provided *in trans* from a cell line. **(d)** Strand specific qRT-PCR recoded IAV segements. **(Top)** Primers targeting GFP coding region. **(Bottom)** Primers targeting lysostaphin coding region. N=3.

Once attached to the cell, the viron is pulled into the cellular endosome where the M2 ion channel reacts to the low pH and releases its genetic cargo, namely eight negative-strand RNA genomic segments, stored as ribonucleoproteins (RNPs) clustered in a 7+1 arrangement (5). These eight RNA molecules encode twelve proteins essential for the IAV lifecycle (PB2, PB1, PB1-N40, PB1-F2, PA, HA, NP, NA, M1, M2, NS1, and NS2). The RNPs are all 13 nm in diameter but range from 30-130 nm in length depending on which segment they include (6, 7). Each RNP is an RNA loop capped by the heterotrimeric RNA-dependent RNA polymerase (RDRP), while the single-stranded RNA is stabilized by one NP monomer per *∼*24 bases. Unlike many other RNA viruses, the viral genome is transported to the nucleus before initiating replication.

In the nucleus, regulatory sequences unique to each segment are recognized by RDRP and transcribed from the negative sense vRNA(-) to positive sense mRNA(+) (Figure 1a). The mRNA(+) is exported to the cytoplasm and translated to generate new viral proteins. Replication of vRNA(-) occurs through a positive sense cRNA(+) intermediate (Figure 1a). During packaging of the virus, one copy of each segment is packaged into each particle in a highly selective manner that is mediated by segment specific 5^*′*^ and 3^*′*^ non-coding sequences that flank each vRNA(-) (7, 8). Some additional residues necessary for viral packaging have been identified within the coding region of the genome (9, 10). Nonetheless, by adding a few additional bases from the coding regions previous studies were able to reprogram the endogenous HA (or another segment) with an alternative protein sequence (11). In this work, we take advantage of sciIAV for therapeutic, research, and bio-manufacturing applications (Figure 1b). First, we generated sciIAV particles encoding proteins that move beyond reporter systems or vaccine development. Second, we leveraged high-throughput sequence-to-phenotype correlations to better understand the constraints of the genetic design space.

## Methods

### Strains and chemicals

IAV strains were stored at −80 °C. All bacterial strains were stored at −80 °C in 20% glycerol solution. Recombinant DNA research was approved by the University of Minnesota Institutional Biosafety Committee. IAV sequences are from the PR8 strain.

### Plasmids

pDZ-PB1, pDZ-PB2, pDZ-PA, pDZ-NP, and pCAGGS-HA were provided by the Langlois lab (12). pDZNA, pDZ-NS, pDZ-M were generated by amplifying the pDZ-PB1 backbone and using Gibson assembly to insert ds-DNA blocks encoding the appropriate PR8 segments. All A-packaged effector plasmids were generated using an isothermal assembly reaction (NEB M5520AA) (13).

### Cell Culture

The MDCK-HA cell line was provided by the Langlois lab. All cells were maintained at 37 °C and 5% CO_2_.Gibco DMEM with GlutaMax (Gibco 10566-016) was supplemented with 10% Heat inactivated-Fetal Bovine Serum (Gibco 10082-147).

### Rescue and amplification of single-cycle rIAV

One million HEK293t cells were plated in either a 6-well plate or a 3 cm dish one day before transfection. The cells were transfected using the Lipofectamine LTX with PLUS Reagent protocol (Invitrogen 15338-100) with 500 ng of each pDZ plasmid (PB1, PB2, PA, NP, NA, NS, and M), pCAGGS-HA, and the desired HA-packaged effector. Twenty-four hours post transfection, 7.5 *×* 10^5^ MDCK-HA cells were overlayed on the transfected HEK293ts with 1 *µ*g/mL of TPCK-treated trypsin (Sigma-Aldrich T1426-50MG). When approximately 70% of the cells had died, 500 *µ*L of supernatant was transferred to a confluent T-75 flask of MDCK-HA cells in OptiMEM (Gibco 11058-021) media with 1 *µ*g/mL of TPCK-treated trypsin. When approximately 70% of the cells had died, the supernatant was collected and centrifuged at 500 rcf for 5 min. The supernatant was separated into 500 *µ*L aliquots and immediately frozen at −80 C. Library experiments were rescued as described above but with 1000 ng of each plasmid.

### Viral quantification

Viral particles were quantified by a plaque assay adapted from Karakus et al. and Sjaastad et al. (14, 15). Briefly, viral particles were thawed on ice. A serial dilution (10^-1^, 10^-2^, 10^-3^, 10^-4^, 10^-5^, 10^-6^) of the viral particles was set up in PBS (Gibco 10010-023). Media was removed from fully confluent MDCK-HA cells in a 6-well plate. Five hundred *µ*L of each dilution was added to the appropriate well and incubated at 37 °C for 1 hr. Next, the cells were covered with 2 mL of 50% DMEM with 1% agar and 2 *µ*g/mL of TPCK-treated trypsin. The cells were incubated upside down at 37 C and 5% CO_2_.

Once plaques formed, 1 mL of 3.7% formaldehyde (Fisher BP531-25) in PBS was added to each well and incubated for 20 min at room temperature. The formaldehyde and agar were removed and 1 mL of 0.5% crystal violet (Sigma C0775-25G) and 20% methanol (Fisher A454-4) in H_2_O was added for 10 min. The crystal violet solution was removed and plaques were manually counted.

### Intracellular RNA collection for qRT-PCR, and vRNA Sequencing

Immediately before collection RNA lysis buffer was made following the protocol developed by Shatzkes et al. (16). Briefly, a 10 mM Tris pH 7.4 (Roche 10812846001), 0.25% Igepal CA-630 (Sigma I8896-50ML),150 mM NaCl (Macron 7581-06) solution was made in H_2_O. Media was aspirated from the cells in a 96-well plate. Two hundred *µ*L of the lysis buffer was added to the cells and incubated for 20 minutes at room temperature. To collect from 6-well plates or 3 cm plates, 1 mL of lysis buffer was used.

### Quantitative Reverse Transcriptase PCR

Intracellular RNA was collected as above. The LunaScript Primer-Free RT Master Mix Kit (NEB E3025S) was used to generate cDNA per manufacturer protocol. Primers were used as listed in Supplementary Note 20. GAPDH primers were adapted from (17). qPCR reactions were set up using the Luna Universal qPCR Master Mix (NEB M3003S). Reactions included 6 *µ*L of Luna Mastermix, 0.05 *µ*L of each primer, 0.6 *µ*L of reverse transcripase mix, 3 *µ*L of RNA, and 2.3 *µ*L H_2_O and run according to manufacturer protocol.

ΔCq values were calculated as the RNA Cq - GAPDH Cq. Normalized expression was calculated as 2^-ΔCq^. Significance was calculated with the two-sided Student’s t-test with Bonferroni correction. Included one biological replicate and three technical replicates per condition.

### Viral particle RNA collection

RNA from viral particles was collected using the Zymo Quick-DNA/RNA Viral MagBead Kit (Zymo R2141) according to manufacturer protocol.

### Reverse transcription for sequencing

LunaScript Primer-Free RT Master Mix Kit (NEB E3025S) was used to generate cDNA using primers oAS344 for cRNA and oAS345 for vRNA (Supplementary Note 21).

Primers oAS344 and oAS345 were then both used to amplify the cDNA using CloneAmp PCR Master Mix (Takara 639298) according to manufacturing protocol. Amplicons were sent for sequencing with Plasmidsaurus. Sequencing was completed on one biological replicate per strain.

### Growth inhibition assay

HEK293ts were grown to *∼* 70% confluence and media was changed to OptiMEM. The cells were transduced with *∼* 1 PFU/cell of either GFP-sciIAV or Lyso-sciIAV and incubated for 24 hrs. *S. aureus* MW2 was streaked on an LB agar plate with 100 *µ*g/mL Ampicillin and grown overnight at 37 °C. A single colony was picked and grown in LB broth with 100 *µ*g/mL Ampicillin at 37 °C in a shaker incubator. Two mL of culture was spread over LB agar plates with 100 *µ*g/mL Ampicillin and 5% defibrinated sheep’s blood (Hardy Diagnostics SB30) and allowed to rest for 2 min. Supernatant was removed and remaining moisture allowed to dry.

Supernatant from transduced HEK293ts was collected and diluted in PBS. Five *µ*L of each dilution was spotted in six replicates and allowed to dry. The plates were incubated overnight at 37 °C.

Zones of inhibition were quantified using ImageJ to count pixels in the zone of inhibition and converted to area. Significance was calculated using a one-way ANOVA. Inhibition assay was done on two biological replicates and three technical replicates per condition.

### Contamination clearance assay

Approximately 25,000 HEK293t cells were plated in a 96-well plate and transduced with either GFP-sciIAV or Lyso-sciIAV at differing MOI (PFU/cell). In the meantime, *S. aureus* RN4220 was grown in LB broth at 37 °C. After 24 hrs, 2 *µ*L of *S. aureus* RN4220 (OD600: 2.606 A.U.), was added to each well. After an additional 24 hrs, OD600 readings and GFP fluorescence measurements (Excitation 488 nm, Emission 507 nm) were taken of each well. One-way ANOVA was used to determine significance within groups indicated in Figure 2c. We fit the data to the Hill Equation using the Gauss-Newton algorithm. Clearance assay was done on two biological replicates and three technical replicates per condition.

**Fig. 2.**
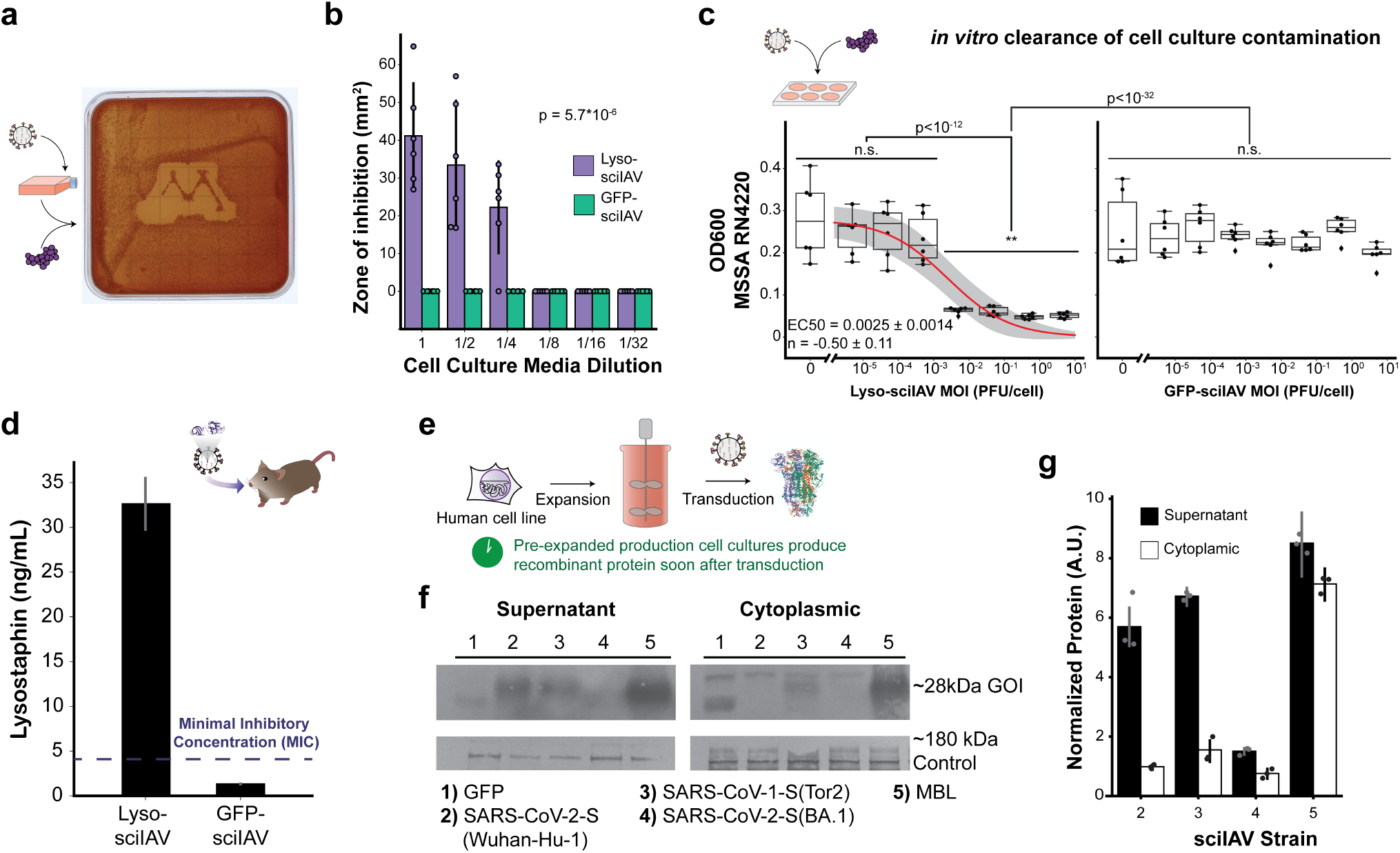
Production of proteins of therapeutic and biomanufacturing interest using domesticated IAV. **(a)** Supernatant from HEK293t cells transduced with lyso-sciIAV inhibits growth of *S. aureus* MW2. **(b)** Zone of inhibition generated by supernatant dilutions from transduced HEK293t cells with measured area. N=6 **(c)** OD600 readings of HEK293t cells transduced with sciIAV strains and contaminated with *S. aureus* RN4220. N=6. **(d)** Production of lysostaphin in transduced C57BL/6 mouse lungs. N=3. **(e)** Schematic of IAV biomanufacturing pipeline. **(f)** Western blot of proteins manufactured using sciIAV platform. **top** Gene of Interest protein products *∼*28 kDa. **bottom** *∼*180 kDa off-target anti-His-antibody band. **(g)** Quantification of protein gel in (f). GFP titer was not plotted since the staining by the anti-His antibody was serendipitous and not likely to occur at a comparable molar efficiency with the other proteins. Protein production was normalized to the *∼*180 kDa band. N=3.

### Mouse treatment and ELISA

C57BL/6 mice were given 10^5^ PFU of either lyso-sciIAV or GFP-sciIAV intranasally. Mice were sacrificed and lung tissue removed after 24 hrs. Protein was collected by sonication. NUNC-Immuno 96-well plates (SIGMA-M9410-1CS) were coated with 100 *µ*L of 4 *µ*g/mL polyclonal anti-lysostaphin antibody (AICBIOTECH-PAb102) in 1*×*ELISA coating buffer (BIOLEGEND-421701). Wells were washed with 0.05% Tween 20 (SIGMA-P9416-50ML) in PBS, blocked with 200 *µ*L of 3% BSA in PBS, and washed again. One hundred *µ*L of protein samples or lysostaphin standards (SIGMA-L9043-5MG) diluted in 0.01% Tween 80 (SIGMA-P5188-100ML) and 0.1% BSA in PBS were added per well and incubated at 37 °C for 1 hr. The polyclonal anti-lysostaphin antibody (AICBIOTECH-PAb102) was conjugated to horseradish peroxidase per the Lightning-Link HRP Antibody Labeling Kit (NOVUS BIOLOGICALS-701-0030) protocol. The plate was washed again and 100 *µ*L of the conjugate was added and incubated at 37 °C for 1 hr. After washing the plate, 100 *µ*L of TMB reagent (SIGMA-RABTMB3-12ML) was added and a kinetic curve measured at 652 nm and 370 nm on a plate reader. Samples were done in biological and technical triplicate.

Animal experiments were approved by the Institutional Animal Care and Use Committee (IACUC) at the University of Minnesota.

### Protein gel and quantification

HEK293t cells were transduced with *∼* 0.5 PFU/cell with each sciIAV strain. After 24 hrs supernatant was collected and concentrated by the Vivaspin-6 5 kDa MWCO filters (Sartorius VS06S1). Cells were lysed in 20 mM Tris-HCl 7.4 pH at 4 °C for 1 hr. The lysed cells were spun down at 17,900 rcf for 20 min. Protein from both fractions was incubated with *β*-mercapethanol (BioRad 161-0710) according to 4*×*Laemmli Sample Buffer (BioRad 161-0747) protocol. Next, 10% Mini-PROTEAN TGX Precast Protein Gels (BioRad 4561033) were loaded with sample and with ladder (Abcam ab116027).

Nitrocellulose transfers were stained with anti-His (Invitrogen PA1-983B), anti-SARS-CoV-S (Absolute Antibodies CR3022), and anti-MBL (SDIX 2445.00.02) antibodies at 1 *µ*g/mL. Secondary staining was done with an antiRabbit antibody conjugated to horseradish peroxidase (Invitrogen G21234) at 1 *µ*g/mL. The gel was visualized using Thermo Scientific Pierce 1-Step Ultra TMB-Blotting Solution (Thermo 37574).

### Library generation

Each library was generated by amplifying HA-packaged GFP rescue plasmid with primers in Supplementary Note 22, digesting with *Dpn*I (NEB R0176L), and assembling by isothermal assembly. Bacteria were electroporated and the full transformation was grown in 2 L LB. The plasmid library was collected by Machery-Nagel Nucleobond Xtra Maxi Prep Kit (Machery-Nagel 740416.50). The library was rescued and expanded as described above.

### Library sorting and RNA extraction

Supernatant was collected after initial rescue in HEK293t cells and from MDCK-HA cells used to backpassage for the packaging and propagation libraries respectively. vRNA was collected by the Zymo Quick-DNA/RNA Viral MagBead Kit (Zymo R2141) according to manufacturer protocol.

HEK293t cells were transduced with the library at *∼*0.4 MOI. After 24 hrs the cells were collected and fluorescence-activated cell sorting (FACS) was done at the University of Minnesota University Flow Cytometry Resource using a BD FACSAria II (P69500132). Collected populations were spun down at 500 rcf for 5 min. Supernatant was removed and intracellular RNA was collected using the NucleoSpin RNA Plus XS kit (Machery-Nagel 740990.250).

### Library sequencing

cDNA of each library was generated using oAS345 (Supplemtary Note 21) and LunaScript Primer-Free RT Master Mix Kit (NEB E3025S).

cDNA and the plasmid library were amplifed using CloneAmp PCR Master Mix (Takara 639298) and primers for amplicon sequencing (Supplemtary Note 23). The amplicons were sent to Genewiz for sequencing.

The Zipf-Mandelbrot distribution, 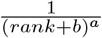, for the 3^*′*^ library was fit using the Gauss-Newton algorithm. Goodness of fit was determined using the Kolmogorov–Smirnov test. Library experiments were done in one replicate each.

## Results and Discussion

### Recoding sciIAV for antibiotic production

New IAV particles can be rescued in cell culture using a pool of 8 (one for each segment) dual expression plasmids that express the vRNA(-) via RNA polymerase I promoters and mRNA(+) via RNA polymerase II promoters (12, 18). We express a recombinant HA vRNA(-) encoding a protein of interest instead of the endogenous product and separately provide HA encoding mRNA(+) (Figure 1c). New viral particles encode our desired protein but, without packaged HA-encoding RNAs, are unable to produce infectious progeny particles. This strategy has been previously used to encode reporters or develop bivalent vaccines (11).

We generated sciIAV strains, expressing either GFP (GFP-sciIAV) or lysostaphin (lyso-sciIAV), a metalloprotease that targets penta-glycine bridges present in the peptidoglycan layer of *Staphylococcus spp*. bacteria (19, 20). Previous studies of IAV reported “superpromoter” mutations in the 5^*′*^ regulatory region of the vRNA(-) that increase the production and expression of the mutant segment (21). We generated additional GFP and lysostaphin strains with these mutation (spGFP-sciIAV and spLyso-sciIAV respectively).

We confirm the presence of both (-) and (+) strand RNA species in a range of transduced cells. HEK293t, A549, and wild-type MDCK (MDCK-Wt) cells were transduced with sciIAV carrying either GFP or lysostaphin. After 24 hours, RNA from lysed cells was used to generate cDNA using either cRNA(+)/mRNA(+) or vRNA(-) specific primers. qRT-PCR confirmed high levels of both strands (Figure 1d) in a strain specific manner. We used RT-PCR with primers targeting the HA regulatory region to amplify the entire fragment for sequencing confirmation (Supplementary Note 1). Noting that the superpromoter variants performed worse than the natural variant, the remainder of our work focused on the reference PR8 regulatory sequences.

### Lysostaphin encoding sciIAV generates bacteria-inhibiting supernatant in cultured cells

We wanted to explore potential therapeutic applications of sciIAV by recoding it to produce an antibacterial protein. Lysostaphin has attracted attention for its potential to combat drug resistant strains of *Staphylococcus aureus* (20, 22–26). Unfortunately, delivery to infected tissues and bioavailability remain barriers to its application. We sought to take advantage of the transient genetic delivery afforded by the sciIAV vector system to bring lysostaphin production directly to the site of infection. We initially characterize the antimicrobial potential of lysosciIAV through a growth inhibition assay. We transduced 3 *×*10^6^ HEK293t cells with 1 MOI of either lyso-sciIAV or GFP-sciIAV. Supernatant from lyso-sciIAV transduced cell lines created a zone of inhibition when spotted onto LB blood agar plates coated with *S. aureus* MW2, a high-virulence community-acquired MRSA strain, n=6 (Figure 2a-b, Supplementary Note 2) (24–26).

### Low-dose protection of cell culture from bacterial infection

We next sought to demonstrate protective activity of lyso-sciIAV in culture. We transduced HEK293t cells with lyso-sciIAV across a range of concentrations from 5 *×*10^-6^ to 5*×*10^0^ MOI of lyso-sciIAV. At 24 hours, we directly contaminated the cell culture with *S. aureus* RN4220 liquid culture (Figure 2c, Supplementary Note 3). Addition of virus did not elicit any significant difference on absorbance in the absence of bacterial contamination (Supplementary Note 3). In contaminated samples, however, lyso-sciIAV was able to completely inhibit the growth of *S. aureus* with doses as low as 0.005 MOI. The reported minimal inhibitory concentration for lysostaphin is reported as 0.004 *µ*g/mL (27), implying that lysostaphin is produced at an estimated 1 pg per infected cell. We calculated an EC50 of 0.0025*±*0.0014 MOI and a Hill coefficient of −0.50*±*0.11.

GFP-sciIAV did not produce any inhibition, indicating that cellular antiviral response did not generate a concurrent antimicrobial response. Interestingly, GFP fluorescence was below the detection limit of our plate reader at 0.005 MOI GFP-sciIAV transduction, speaking to the potency of lysostaphin (Supplementary Note 4).

### Therapeutically relevant production titers of lysostaphin in mouse lung tissue

A major promise of our transient, non-integrating sciIAV platform is the potential for delivery of antimicrobial proteins to lung tissue as a prophylactic or post-exposure therapy. To ensure that delivery could occur *in vivo*, we treated C57BL/6 mice with 10^5^ PFU of either lyso-sciIAV or GFP-sciIAV. We found that lyso-sciIAV-infected mice generated 32.6 *±* 3.4 ng/mL of lysostaphin, more than 8-fold the minimal inhibitory concentration (Figure 2d) (27). The mice exhibited no adverse symptoms from the treatment. Mice are not natural hosts for clinical *S. aureus*. The introduced bacteria are readily cleared at all but extreme doses, making them a poor model for staphyloccocal pneumonia (28–30). We did not explore the ability of lyso-sciIAV to clear a lung infection in a suitable model such as rabbits.

### Separating cell growth and production in biomanufacturing workflows

In an alternative use-case for domesticated sciIAV, we chose to take advantage of the binary reprogramming exhibited by the sciIAV system for developing an improved biomanufacturing workflow. Traditional mammalian biomanufacturing pipelines begin with engineering a cell line to produce the therapeutic protein. Highly productive lines are selected and then scaled to biomanufacturing volumes, a process that can take weeks. If there is a fitness cost to the productive lines, this can take even longer (5).

We envisioned replacing this slow, linear workflow with a parallelized manufacturing paradigm. Wild-type cells would be grown to production culture volumes, avoiding fitness costs often associated with recombinant cell lines. In parallel, recombinant sciIAV particles are created and expanded. The production culture is then transduced at ≫1 MOI, rapidly reprogramming nearly all of the cells from no production to high production (Figure 2e, Supplementary Note 5). Such a workflow takes advantage of the rapid sciIAV expansion rate compared to that of mammalian cells, reaching a sufficient titer of sciIAV particles in a few days to infect cultured cells that had been expanding for several weeks. Such a workflow is particularly attractive when producing a protein of interest as quickly as possible is important. For example, evolution of new SARS-CoV-2 variants during the COVID-19 pandemic required routine at-scale production of Spike (S) protein receptor binding domain variants for testing commercial antigen assays.

We designed and rescued sciIAV strains encoding the Spike protein receptor binding domain (SPRBD) from various SARS-CoV strains. Note that we are not expressing the entire spike protein, only the C-terminal fragment that resides outside of the viral particle. This work was approved by the UMN Institutional Biosafety Committee and does not present a risk of changing the tropism or infectivity of the sciIAV particles. Specifically, we designed sciIAV particles to express SARS-CoV-2 (reference strain, Wuhan-Hu-1) SPRBD, SARS-CoV-1 (reference strain, Tor2) SPRBD, SARS-CoV-2 (BA.1) SPRBD, and Mannan Binding Lectin (MBL). Each of these included a signal sequence for secretion. We were able to rescue and confirm sequencing of each of these strains (Supplementary Notes 6, 7, 8, 9).

We collected protein from cells transduced with 0.5 MOI with these recombinant sciIAV strains. The anti-His antibodies used in our Western Blot had large off-target binding but were able to reveal our protein targets (Figure 2f, g, Supplementary Note 10). The GFP control did not include a signal sequence. Though some protein was detected in the culture supernatant, the majority of the product remained cytoplasmic. Notably the GFP we expressed did not include a 6 *×* His-tag so co-staining comes from off-target binding. While the SARS-CoV-2 (Wuhan-Hu-1) SPRBD, SARS-CoV-1 SPRBD (Tor2), and MBL exhibited expression, the BA.1 variant would not express across multiple attempts.

Furthermore, MBL signal was robust in both supernatant and cytoplasm indicating its expression was much higher than any of the SARS-CoV SPRBD variants. This highly variant protein production was seen despite using identical regulatory sequences outside of the coding RNA sequence, indicating the presence of compositional genetic context effects (31).

### Interrogating cryptic regulatory regions in the protein coding sequence of IAV HA Segment

The irregularity of expression between (even similar) coding sequences led us to question the independence of the regulatory and coding regions. To test for potential genetic context dependence, we adapted the sciIAV platform for high-throughput screening of thousands of IAV variants for cryptic regulatory elements.

We based our screen on the GFP-sciIAV reporter as this cargo expresses well and GFP is known to tolerate a broad range of N-terminal additions without loss of fluorescence (32–34). We used degenerate oligos to add 45 randomized bases (15 randomized amino acids/stop codons) to the N-terminus of GFP immediately following the ATG start codon. We transduced HEK293t cells at *∼* 0.4 MOI with the rescued sciIAV library and used FACS to collect high-, medium-, low-, and non-GFP cells for next-generation sequencing (Figure 3a).

**Fig. 3.**
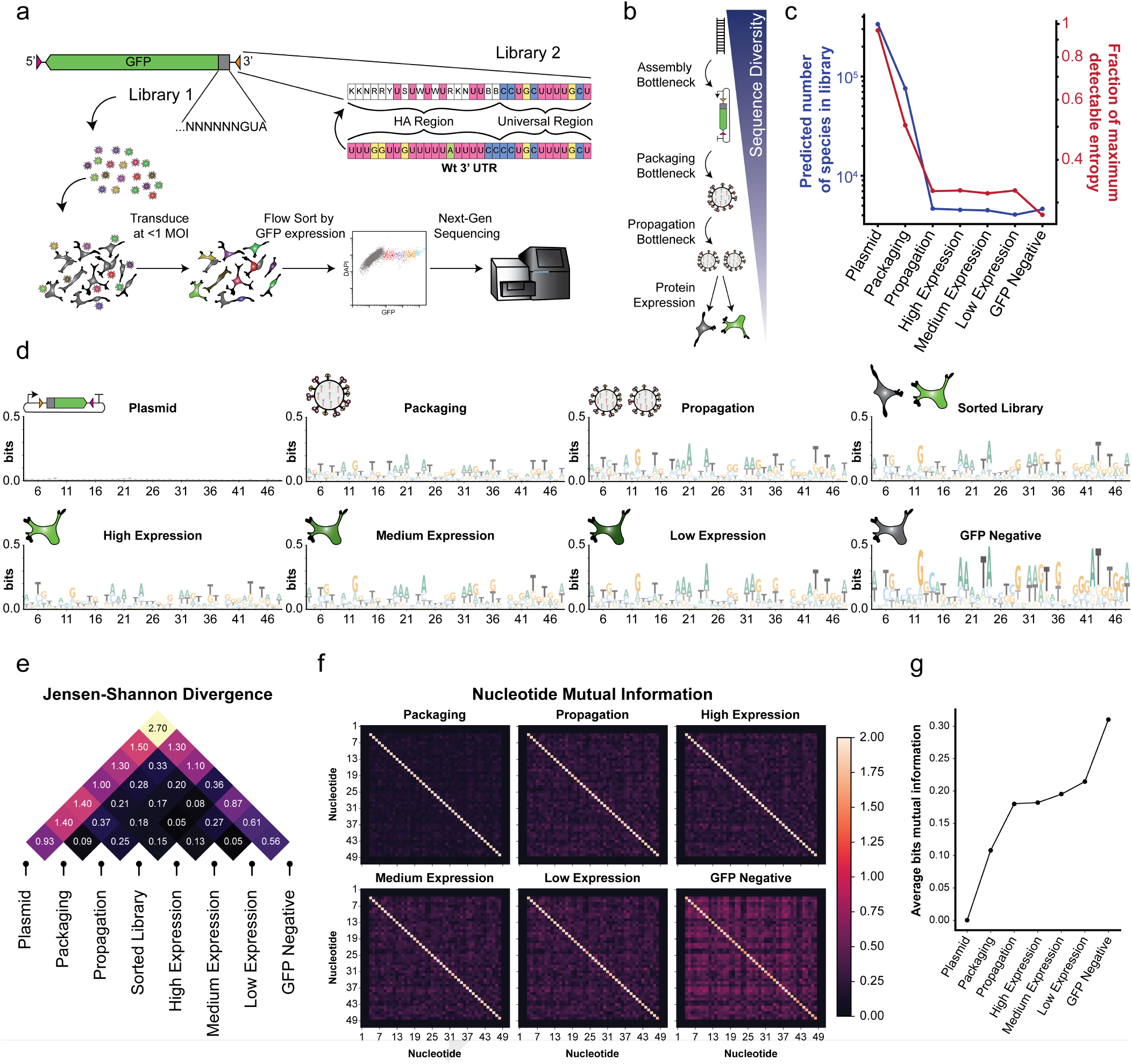
High-throughput screening of genetic context effects on IAV protein expression. **(a)** High-throughput library generation and screening pipeline. **(b)** Schematic of major bottlenecks in IAV library generation and expression selection. **(c)** Library diversity and species representation over selection bottlenecks and across GFP expression levels. **(d)** Sequence logos for each library. **(e)** Quantified divergence between libraries using the sum of basepair level Jensen-Shannon Divergence with higher values indicated larger difference between libraries. **(f)** Pairwise nucleotide level mutual information. **(g)** Average mutual information over selection bottlenecks and across GFP expression levels.

We estimated the total number of unique species at each phase of selection using ecological modeling methods (35). The packaging and propagation bottlenecks represented large losses in species representation. However, there were no major losses in number of species lost after fluorescence level selection (i.e. each population had the same number of species as the propagated viral library within a 95% confidence interval) (Figure 3b and c). As a measure of diversity, we tracked the total entropy at each stage normalizing to the highest entropy we could theoretically detect. Despite the number of species staying the same across the last samples, there was a 5% fractional entropy drop in the GFP-negative cells (Figure 3c).

The plasmid library had effectively maximal entropy at every base and amino acid in the library (Figure 3c, d, Supplementary Note 11). Meanwhile, pairwise mutual information was minimized between each nucleotide (Supplementary Note 11). These indicate no bias was introduced during the cloning of rescue plasmids.

We generated sequence logos to represent the distribution of sequences at each step (Figure 3d). We processed the high-, medium-, low-, and non-GFP populations separately but also analyzed them as one whole sequence library. JensenShannon Divergence (JSD) at each base position was used to map enrichment between sequenced libraries (Supplementary Note 12) (36). JSD is a measure of difference in probability distribution at each base ranging 0≤ JSD≤ 1, with higher JSD representing more similar distributions. We summarized the differences by summing the JSD across all bases (Figure 3e). We similarly analyzed the first 48 bases of all HA sequences available on GenBank.

Packaging led to weak enrichment of a single nucleotide at most base coordinates (Figure 3d, Supplementary Note 13). This enrichment was chiefly replicated but slightly exaggerated in the propagation step (Figure 3d, Supplementary Note 14). The propagation bottleneck selection could occur by three main mechanisms: differential packaging efficiency, differential replication by the RDRP, or differential life cycle length. The low propagation vs. packaging JSD indicates the selection mechanisms are similar between the two bottlenecks (Figure 3e, Supplementary Note 12).

We compared the enrichment after the packaging and propagation bottlenecks to the enrichment present in known HA sequences. At 15 of the 45 bases in the library, the enriched nucleotide matched that of known HAs. More than half of these were thymines (i.e. uracils in the viral life cycle). Most of these relatively conserved residues were in the first position for codons within the +1 reading frame. The whole library after FACS diverged from the viral library used to tranduce the HEK293ts despite the high similarity during propagation (Figure 3e, Supplementary Note 12).

After FACS, the JSDs for each population trended lower for populations with more similar expression indicating the assay was able to capture differences in expression. The high-, medium-, and low-expression populations were nonetheless relatively similar. Strikingly, the negative expression population had large JSDs when compared to the other populations or any of the library generation steps (Figure 3d, e, Supplementary Notes 15, 16, 17, 18, 19). This suggests that rather than requiring or preferring certain sequences, IAV protein expression is intolerant of certain sequences.

We next looked at the basepair mutual information, a measure of pairwise correlation, through the assay (Figure 3f, g). It was immediately clear that lower expressing populations had higher mutual information. We investigated if this was due to secondary structure by comparing expression to folding energy (Δ*G*), however found no correlation (R^2^ = −0.01). The global increase in mutual information then indicates selection for a particular sequence or small set of sequences, consistent the idea that the difference in expression is driven by intolerance of some sequences.

### Library based screening of 3^*′*^ UTR sequence constraints

The 3^*′*^ UTR of each IAV segment shares a short universal region followed by a region specific to each segment. We sought to interrogate the sequence restrictions on the HA segment specific region through a similar library method (Figure 3a). Since we were modifying the canonical regulatory sequences, we expected a stringent selective pressure on this library. To overcome this barrier we used a more limited library informed by predicted secondary structures (Supplementary Note 24)(37). We focused on mutations that would most likely impact folding (positively or negatively) and those that would impact the Kozak sequence. Our library was sized to ensure high probability that the wildtype sequence would be present in the library as an internal control.

The parental sciIAV-GFP strain retains sequence from HA on the 3^*′*^ end of the GFP CDS as a way to boost expression of the cargo gene (9–11). In our library, we excluded this region to focus on just the impact of the UTR itself. As a corollary outcome, this change in sequence length enables discrimination between the wildtype UTR generated in our library and the parental sequence from which the library was created (70 bp and 144 bp respectively)(Figure 4a).

**Fig. 4.**
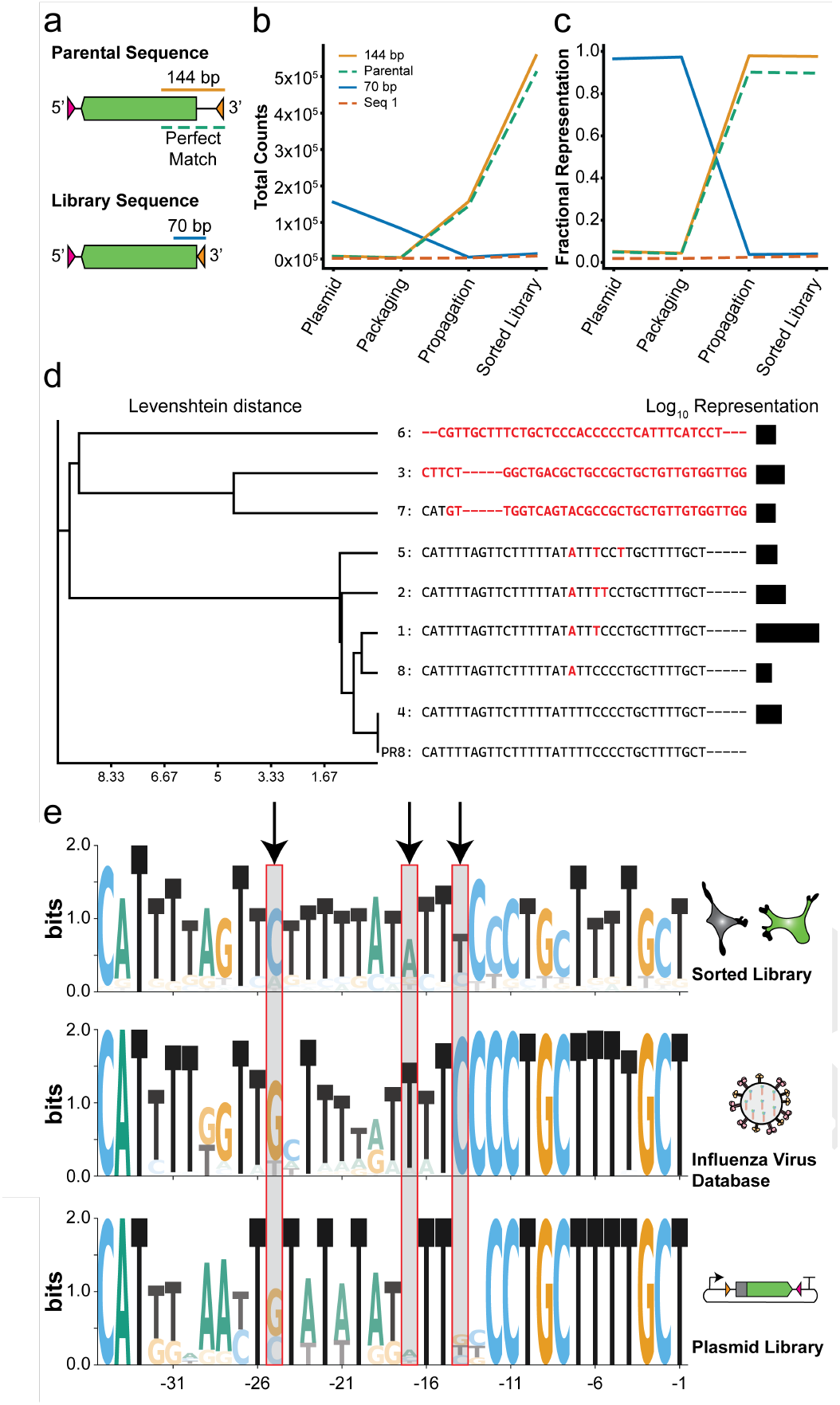
High-throughput analysis of 3^*′*^ UTR sequence restrictions. **(a)** Schematic comparing the genetic structure and differences between parental plasmid used to template library and the library sequence. **(b)** Total counts of populations within the sequenced library at individual experimental stages. **(c)** Relative representation of populations within the sequenced library at individual experimental stages **(d)** Levenshein distance based phylogeny of primary sequences present in sorted library, aligned sequences, and relative representation within the library.**(e)** Sequence logos generated from unique sequences present in either the sorted library, plasmid library, or reported sequences from the Influenza Virus Database.

As for the previous library, we measured the sequence diversity in our library at each step in the experiment (Figure 4b). Despite *Dpn*I treatment aimed to eliminate the parental sequence after the initial cloning steps, a small minority of parental plasmid remained in the library. It represented *∼* 3.4% of the total reads at the plasmid stage. Breaking these parental reads down further, 3.1% were a perfect match to the reference sequence, and 0.3% were single nucleotide polymorphisms (SNPs) of the parental seqeunce, which could either represent sequencing artifacts or rare mutations that occured during plasmid propogation. This low ratio of SNPs to perfectly aligning reads suggests that sequencing artifacts represent a low fraction of the diversity throughout the entire library.

Approximately 97% of the reads at the plasmid stage aligned to the 70 bp UTR region expected in our diversity library. They followed a Zipf-Mandelbrot distribution (Supplementary Note 25), with singletons accounting for 25% of the reads (representing 50% of the sequence diversity), doublets accounting for 11% of the reads (22% of the sequence diversity), triplets accounting for 5% of the reads (11% of the sequence diversity), et cetera.

The 144 bp parental population remained a minority in the packaging library ( ∼2.6%). However, in the propagation and sorted libraries, the representation shifted to the parental population representing ∼98% of the total (Figure 4b and c). This suggests a strong fitness advantage for the wild-type UTR sequence. Interestingly, we could detect a similar strong positive selection for the wild-type UTR sequence that was generated in our diversity library (70 bp UTR sequences). This sequence was enriched from undetectable amounts in the plasmid stage to a substantial fraction of the 70 bp UTR sequences at the packaging and propagation stages.

By the sorted library only 8 sequences were present with a read depth greater than ten. We aligned this group and used the Levenshtein distance metric to cluster them (Figure 4d). As expected, most sequences from the library were de-enriched over the course of the experiment. At all but two nucleotides (HA 17 and 19 in 3^*′*^-5^*′*^ vRNA coordinates) the library enriched primarily towards the PR8 sequence. Among these, the PR8 reference sequence was regenerated indicating the library selection gave trustworthy results. Sequences 1, 2, 4, 5, and 8 approximated the PR8 sequence and shared a U *⇒* A change expected to increase the size of an interior loop in the vRNA (Supplementary Note 24). It is less clear what the impact may be of the other changes on secondary structure. Sequences 3, 6, and 7 do not resemble the PR8 sequence at all and do not include even the universal region of 3^*′*^ UTR.

We collected each unique sequence that appeared in the selected library as well as the plasmid library, generating a sequence logo (Figure 4e). We chose not to bias according to sequence frequency to focus on possible rather than dominating sequences. We then collected the HA segment sequences available on Influenza Virus Database and analyzed similarly (38). For the Influenza Virus Database data we excluded any sequence with ambiguous bases reported in the 3^*′*^ UTR. Again the library largely suggested selective pressure to maintain wild-type sequence with HA 17 and 19 being the primary exceptions. PR8 differs from the primary G at HA 25 and this difference is reflected in our library as well.

Based on comparison of GFP expression levels, the IAV experienced a stronger selection within the GFP negative population. Indeed, a general trend was clear where bases common in the GFP negative population were progressively rarer in higher expressing populations. These data suggest that our assay was better at identifying poorly expressed sequences than highly expressed sequences. Folding Δ*G* does not correlate with expression, indicating that secondary structure is unlikely to be causing the inhibited expression. Instead, given that the predicted number of species was approximately consistent between populations, it is likely that multiple discrete motifs are driving the difference. The relatively global increase in mutual information corroborates this as base pairing would present was individual points. It is likely that several, mutually exclusive motifs in the library can decrease protein production. There appears to be at least some tolerance to every nucleotide at every position supporting the idea that multiple discrete motifs drive the effect.

As expected the 3^*′*^ UTR library had more stringent sequence requirements than the N-terminus library. Notably, the library recaptured the wild-type sequence. Five of the sequences we identified shared high similarity to the wild-type sequence, differing by up to three nucleotides. The U ⇒A change at the nineteenth position was present in four of the five sequences even though the plasmid level library contained all four nucleotides.

In future work, individual sequences identified in this experiment can be rescued and tested based on their similarity to the consensus. This would allow further investigation of how mutating specific nucleotides impacts the expression and help identify the core motifs. In the meantime, the library generation approach can be used to probe other parts of the IAV genome. The specific design we used would be best for probing the UTRs, N-terminus, and C-terminus of any segment. Other portions of GFP known to be amenable to modification are also potential targets.

Identification of specific motifs moving forward will improve the ability to bypass genetic context restrictions of the IAV gene delivery system. This work did not investigate mechanistic explanations for the outcome. RDRP or NP binding preferences may contribute. Alternatively, host cell factors may interact with the sequences. Domestication efforts moving forward may seek to eliminate or direct the context effects through rational engineering or directed evolution.

This work takes a domestication approach toward utilizing the properties of IAV for therapeutics, biomanufacturing, and research. We develop a strain capable of producing inhibitory levels of an antibiotic protein *in vitro* and *in vivo*. We then adapt the platform for a biomanufacturing pipeline. Finally, we study genetic context effects on protein expression in the HA segment identifying potential future directions of study.

## Data Availability

The data and materials that support the results or analyses presented in this paper are freely available upon request.

## A. Acknowledgements

AS was supported by a Biotechnology Training Grant: NIH T32GM008347. CSS, ANH, and MJS were partially supported by start-up funds awarded by the University of Minnesota.

## Contributions

CSS and MJS conceived the study. AS, CSS, and SNR designed and executed experiments. AS, CSS, and ANH constructed genetic reagents and strains for the experiments. RAL and MJS oversaw the data collection. AS and MJS wrote the manuscript. All authors reviewed and revised the manuscript.

## Bibliography

1. RWH Ruigrok, PJ Andree, RAM Hooft Van Huysduynen, and JE Mellema. Characterization of three highly purified influenza virus strains by electron microscopy. Journal of General Virology, 65:799–802, 4 1984. ISSN 0022-1317. doi: 10.1099/0022-1317-65-4-799.

2. Centers for Disease Control and Prevention. Disease burden of flu, 10 2022.

3. M Yamaguchi, R Danev, K Nishiyama, K Sugawara, and K Nagayama. Zernike phase contrast electron microscopy of ice-embedded influenza a virus. Journal of Structural Biology, 162:271–276, 5 2008. ISSN 10478477. doi: 10.1016/j.jsb.2008.01.009.

4. M Luo. Influenza Virus Entry, pages 201–221. Springer US, 2012. ISBN 978-1-4614-0980-9. doi: 10.1007/978-1-4614-0980-9_9.

5. S Wasilewski, LJ Calder, T Grant, and PB Rosenthal. Distribution of surface glycoproteins on influenza a virus determined by electron cryotomography. Vaccine, 30:7368–7373, 12 2012. ISSN 0264410X. doi: 10.1016/j.vaccine.2012.09.082.

6. T Noda and Y Kawaoka. Structure of influenza virus ribonucleoprotein complexes and their packaging into virions. Reviews in Medical Virology, 20:380–391, 11 2010. ISSN 10529276. doi: 10.1002/rmv.666.

7. AJ Eisfeld, G Neumann, and Y Kawaoka. At the centre: influenza a virus ribonucleoproteins. Nature Reviews Microbiology, 13:28–41, 1 2015. ISSN 1740-1526. doi: 10.1038/nrmicro3367.

8. X Li, M Gu, Q Zheng, R Gao, and X Liu. Packaging signal of influenza a virus. Virology Journal, 18:36, 12 2021. ISSN 1743-422X. doi: 10.1186/s12985-021-01504-4.

9. T Watanabe, S Watanabe, T Noda, Y Fujii, and Y Kawaoka. Exploitation of nucleic acid packaging signals to generate a novel influenza virus-based vector stably expressing two foreign genes. Journal of Virology, 77:10575–10583, 10 2003. ISSN 0022-538X. doi: 10.1128/JVI.77.19.10575-10583.2003.

10. GA Marsh, R Hatami, and P Palese. Specific residues of the influenza a virus hemagglutinin viral rna are important for efficient packaging into budding virions. Journal of Virology, 81: 9727–9736, 9 2007. ISSN 0022-538X. doi: 10.1128/JVI.01144-07.

11. A Nogales, SF Baker, W Domm, and L Martínez-Sobrido. Development and applications of single-cycle infectious influenza a virus (sciiav). Virus Research, 216:26–40, 5 2016. ISSN 01681702. doi: 10.1016/j.virusres.2015.07.013.

12. N Hitoshi, Y Ken-ichi, and M Jun-ichi. Efficient selection for high-expression transfectants with a novel eukaryotic vector. Gene, 108:193–199, 12 1991. ISSN 03781119. doi: 10.1016/0378-1119(91)90434-D.

13. DG Gibson, L Young, R Chuang, JC Venter, CA Hutchison, and HO Smith. Enzymatic assembly of dna molecules up to several hundred kilobases. Nature Methods, 6:343–345, 5 2009. ISSN 1548-7091. doi: 10.1038/nmeth.1318.

14. U Karakus, M Crameri, C Lanz, and E Yanguez. Propagation and Titration of Influenza Viruses, volume 1836, pages 59–88. Springer New York, 2018. ISBN 978-1-4939-8677-4. doi: 10.1007/978-1-4939-8678-1.

15. LE Sjaastad, EJ Fay, JK Fiege, MG Macchietto, IA Stone, MW Markman, S Shen, and RA Langlois. Distinct antiviral signatures revealed by the magnitude and round of influenza virus replication in vivo. Proceedings of the National Academy of Sciences of the United States of America, 115:9610–9615, 9 2018. ISSN 1091-6490. doi: 10.1073/pnas.1807516115.

16. K Shatzkes, Be Teferedegne, and H Murata. A simple, inexpensive method for preparing cell lysates suitable for downstream reverse transcription quantitative pcr. Scientific Reports, 4: 4659, 4 2014. ISSN 2045-2322. doi: 10.1038/srep04659.

17. Y Li, L Sun, W Zheng, M Mahesutihan, J Li, Y Bi, H Wang, W Liu, and TR Luo. Phosphorylation and dephosphorylation of threonine 188 in nucleoprotein is crucial for the replication of influenza a virus. Virology, 520:30–38, 7 2018. ISSN 00426822. doi: 10.1016/j.virol.2018.05.002.

18. M Quinlivan, D Zamarin, Ad García-Sastre, A Cullinane, T Chambers, and P Palese. Attenuation of equine influenza viruses through truncations of the ns1 protein. Journal of Virology, 79:8431–8439, 7 2005. ISSN 0022-538X. doi: 10.1128/JVI.79.13.8431-8439.2005.

19. CA Schindler and VT Schuhardt. Lysostaphin: A new bacteriolytic agent for the staphylococcus. Proceedings of the National Academies of Science, 51:414–421, 3 1964.

20. J Jayakumar, VA Kumar, L Biswas, and R Biswas. Therapeutic applications of lysostaphin against staphylococcus aureus. Journal of Applied Microbiology, 131:1072–1082, 9 2021. ISSN 1364-5072. doi: 10.1111/jam.14985.

21. A Belicha-Villanueva, JR Rodriguez-Madoz, J Maamary, A Baum, D Bernal-Rubio, M Minguito de la Escalera, A Fernandez-Sesma, and Adolfo García-Sastre. Recombinant influenza a viruses with enhanced levels of pb1 and pa viral protein expression. Journal of Virology, 86:5926–5930, 5 2012. ISSN 0022-538X. doi: 10.1128/JVI.06384-11.

22. BN Kreiswirth, S Löfdahl, MJ Betley, M O’Reilly, PM Schlievert, MS Bergdoll, and RP Novick. The toxic shock syndrome exotoxin structural gene is not detectably transmitted by a prophage. Nature, 305:709–712, 10 1983. ISSN 0028-0836. doi: 10.1038/305709a0.

23. D Nair, G Memmi, D Hernandez, J Bard, M Beaume, S Gill, P Francois, and AL Cheung. Whole-genome sequencing of staphylococcus aureus strain rn4220, a key laboratory strain used in virulence research, identifies mutations that affect not only virulence factors but also the fitness of the strain. Journal of Bacteriology, 193:2332–2335, 5 2011. ISSN 0021-9193. doi: 10.1128/JB.00027-11.

24. Centers for Disease Control and Prevention (CDC). Four pediatric deaths from community-acquired methicillin-resistant staphylococcus aureus — minnesota and north dakota, 1997-1999. MMWR. Morbidity and mortality weekly report, 48:707–10, 8 1999. ISSN 0149-2195.

25. T Baba, F Takeuchi, M Kuroda, H Yuzawa, K Aoki, A Oguchi, Y Nagai, N Iwama, K Asano Naimi, H Kuroda, L Cui, K Yamamoto, and K Hiramatsu. Genome and virulence determinants of high virulence community-acquired mrsa. The Lancet, 359:1819–1827, 5 2002. ISSN 01406736. doi: 10.1016/S0140-6736(02)08713-5.

26. C Kim, M Mwangi, M Chung, C Milheirço, H de Lencastre, and A Tomasz. The mechanism of heterogeneous beta-lactam resistance in mrsa: Key role of the stringent stress response. PLoS ONE, 8:e82814, 12 2013. ISSN 1932-6203. doi: 10.1371/journal.pone.0082814.

27. CM Kusuma and JF Kokai-Kun. Comparison of four methods for determining lysostaphin susceptibility of various strains of staphylococcus aureus. Antimicrobial Agents and Chemotherapy, 49:3256–3263, 8 2005. ISSN 0066-4804. doi: 10.1128/AAC.49.8.3256-3263.2005.

28. HK Kim, D Missiakas, and O Schneewind. Mouse models for infectious diseases caused by staphylococcus aureus. Journal of Immunological Methods, 410:88–99, 8 2014. ISSN 00221759. doi: 10.1016/j.jim.2014.04.007.

29. DM Mrochen, LM Fernandes de Oliveira, D Raafat, and S Holtfreter. Staphylococcus aureus host tropism and its implications for murine infection models. International Journal of Molecular Sciences, 21:7061, 9 2020. ISSN 1422-0067. doi: 10.3390/ijms21197061.

30. W, Salgado-Pabón., and PM Schlievert. Models matter: the search for an effective staphylococcus aureus vaccine. Nature Reviews Microbiology, 12:585–591, 8 2014. ISSN 1740-1526. doi: 10.1038/nrmicro3308.

31. S Cardinale and AP Arkin. Contextualizing context for synthetic biology - identifying causes of failure of synthetic biological systems. Biotechnology Journal, 7:856–866, 7 2012. ISSN 18606768. doi: 10.1002/biot.201200085.

32. NM Escobar, S Haupt, G Thow, P Boevink, S Chapman, and K Oparka. High-throughput viral expression of cdna–green fluorescent protein fusions reveals novel subcellular addresses and identifies unique proteins that interact with plasmodesmata. The Plant Cell, 15:1507–1523, 7 2003. ISSN 1040-4651. doi: 10.1105/tpc.013284.

33. A Hayashi, D Da-Qiao, C Tsutsumi, Y Chikashige, H Masuda, T Haraguchi, and Y Hiraoka. Localization of gene products using a chromosomally tagged gfp-fusion library in the fission yeast schizosaccharomyces pombe. Genes to Cells, 14:217–225, 2 2009. ISSN 13569597. doi: 10.1111/j.1365-2443.2008.01264.x.

34. Kazuhide Misawa, Tetsuya Nosaka, Sumiyo Morita, Azusa Kaneko, Tatsutoshi Nakahata, Shigetaka Asano, and Toshio Kitamura. A method to identify cdnas based on localization of green fluorescent protein fusion products. Proceedings of the National Academy of Sciences, 97:3062–3066, 3 2000. ISSN 0027-8424. doi: 10.1073/pnas.97.7.3062.

35. A Chao. Estimating the population size for capture-recapture data with unequal catchability. Biometrics, 43:783, 12 1987. ISSN 0006341X. doi: 10.2307/2531532.

36. M Nettling, H Treutler, J Grau, J Keilwagen, S Posch, and I Grosse. Difflogo: a comparative visualization of sequence motifs. BMC Bioinformatics, 16:387, 12 2015. ISSN 1471-2105. doi: 10.1186/s12859-015-0767-x.

37. M Zuker and P Stiegler. Optimal computer folding of large rna sequences using thermodynamics and auxiliary information. Nucleic Acids Research, 9:133–148, 1981. ISSN 0305-1048. doi: 10.1093/nar/9.1.133.

38. Y Bao, P Bolotov, D Dernovoy, B Kiryutin, L Zaslavsky, T Tatusova, J Ostell, and D Lipman. The influenza virus resource at the national center for biotechnology information. Journal of Virology, 82:596–601, 1 2008. ISSN 0022-538X. doi: 10.1128/JVI.02005-07.

