## SupplementaryInfo for "High-throughput investigation of genetic design constraints in domesticated Influenza A Virus for transient gene delivery"

### Supplementary Note 1: 5′ to 3′ vRNA sequence of lyso-sciIAV

AGTAGAAACAAGGGTGTTTTTCCTCATATTTCTGAAATTCTAATCTCAGATGCATATTCTGCACTGCAAAGAT
CCATTAGAACACATCCAGAAACTGATTGCCCCCAGGGAGACCAAAAGCACCAGTGAACTGGCGACAGTTGAGT
AGATCTCGAGTCACTTAATGGTGCCCCACAGCACTCCCAGTGTGTTGGTGGACTTCTGCCATGTCCGCACAGG
CAGATAGATCCGCTGGCCGCTATTGCCTGTGTATCCGACCCACACGTGGCCGTCCTGCTTCATCACTTCGTCG
TAGTGGATGGTCTGTCCGGCTTTCAGCACTCCGCTCTGAGGCATGGATCTGAAAGGTCCGGTGGTCCGGGTGA
TGATGTCGGTGTTTGGGGTGAAGCTGGCGCTCTCGCTCTTGTACAGGGTGCCGTATTTGTTTGTTTTCCAGCC
GGTGTTAGGTGTGGGTGTAACGGTTCCGCCAGCCTTTCCATAGCCGGCGGACTTCAGGAAAGGCATAGGATCC
TGGGCTGTGCTCTGGCTGAAGCTGTTGACCATCCGCTGGAAGTGCAGATGAGGAGCTGTGCTGTAGCCTGTGC
TGCCGCTCCAGCCAATGATCTGGCCGGCCTTCACGTAGTCGCCCACTTTCACGTTGTACTTGCTCAGGTGCAT
GTACCACTGCCGGTGCACGCCGTCATTCTCGATCAGGCCGATCTGATTGCCTCCGCCGTAATTGGACCAGCCG
GCTTCCACAATTTTGCCGCTGCTGATGGCCTTCACAGGGGTGCCGATGTTCATGAAAAAGTCCACGCCGTAGT
GCATGCCGCCATTGATGCCCAGAGGATAGGGGCCGTAGCCGTAGCCTTTCTTGTAGTTGTTCAGCCACTGAGC
AGAATGTTCGTGTGTAGCGGCGGCTTCGGCCACGGCAATACAGATCAGGGCGAACAGgACCTTCACgCCCATG
GCTAGCCTATACAAATTGTGTCTGCAACTGCAGCTGCAAGTGCACTTAACAGGACCAGTAGGTTTGCCTTCAA
TTTGGTTGTTTTTATTTTCCCCTGCTTTTGCT

DRAFT

### Supplementary Note 2: *S. aureus* MW2 growth inhibition assay

Photos of blood agar growth inhibition plates totaling 6 replicates over two plates.

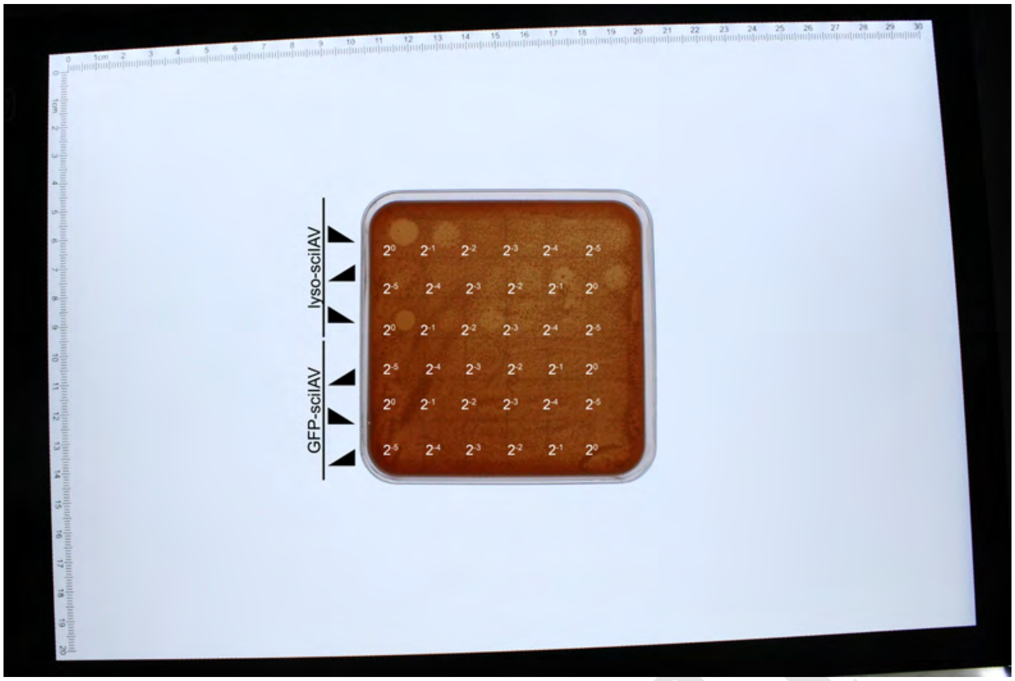


**Fig. 5**

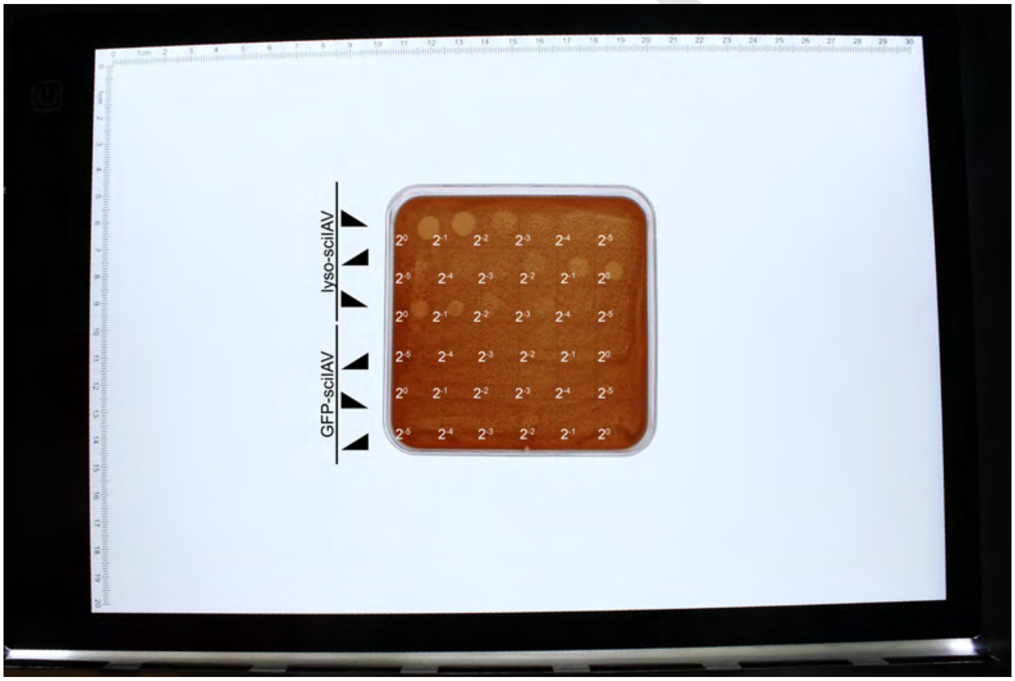


**Fig. 6**

### Supplementary Note 3: *S. aureus* RN4220 cell culture contamination clearance

Optical Density at 600nm measured for cell culture transduced with GFP- or lyso-sciIAV then contaminated with *S. aureus* RN4220. A no contamination control was included.

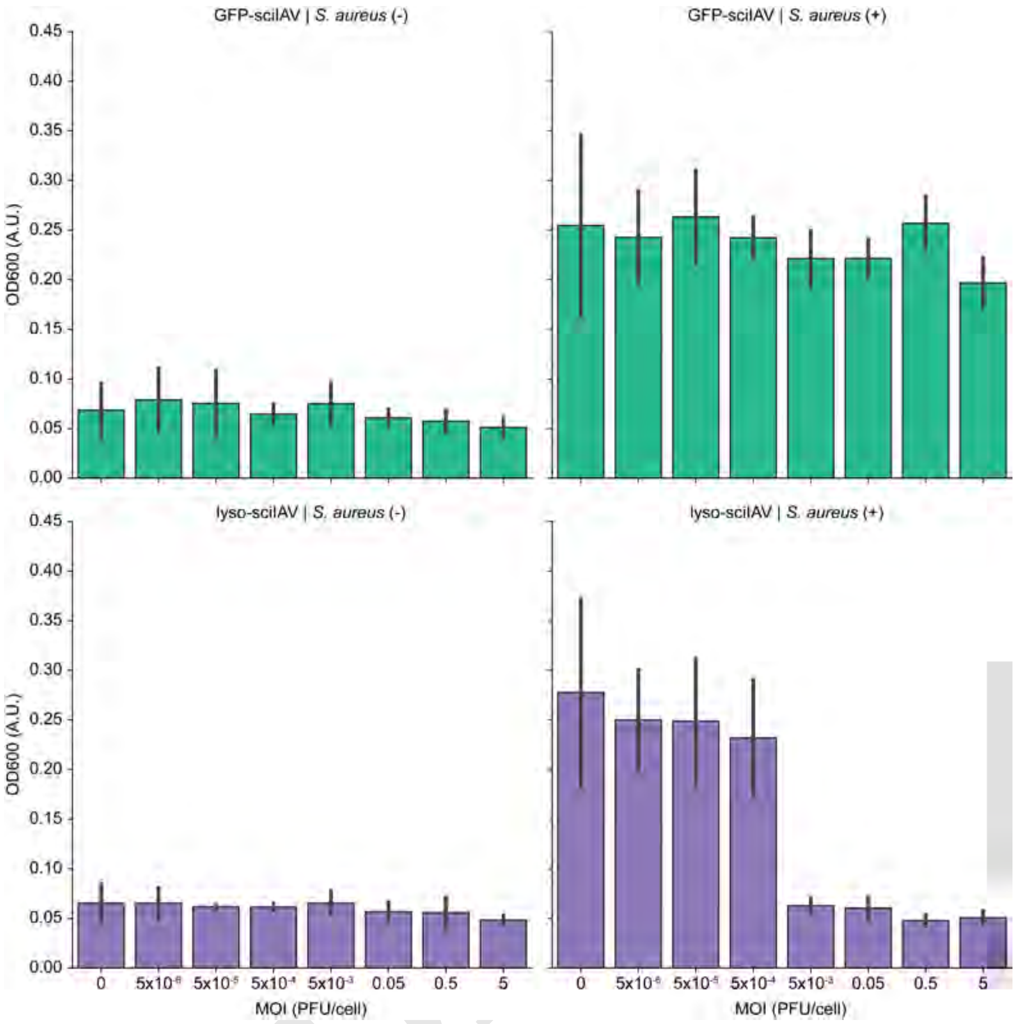


**Fig. 7**

### Supplementary Note 4: *S. aureus* RN4220 cell culture GFP measurement

GFP fluorescence at 600nm measured for cell culture transduced with GFP- or lyso-sciIAV then contaminated with *S. aureus* RN4220. A no contamination control was included.

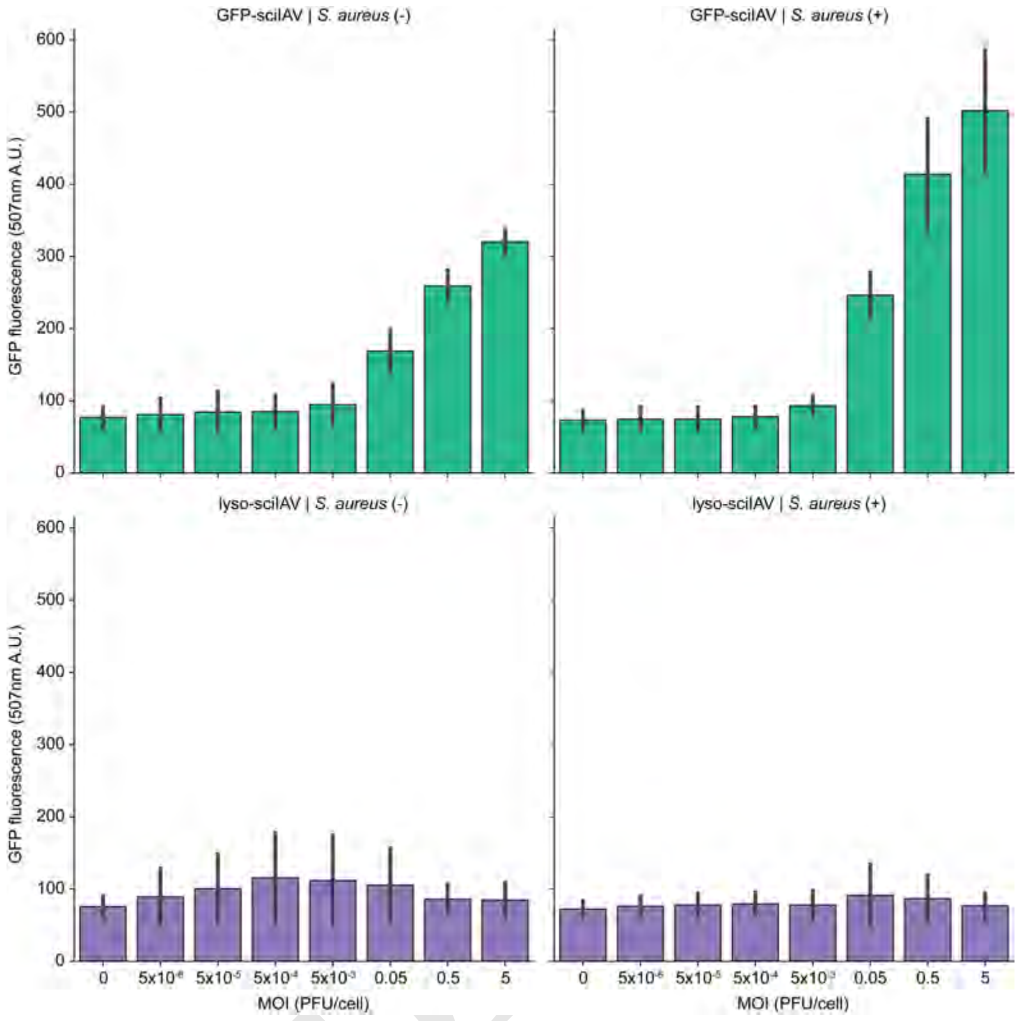


**Fig. 8**

### Supplementary Note 5: sciIAV biomanufacturing platform

Schematic comparing IAV and traditional biomanufacturing. **(Top)** The traditional biomanufacturing pipeline is a linear process. First a cell line must be generated and selected for high production. The best lines are expanded to manufacturing titers taking weeks, extended by any fitness cost posed by the engineering. **(Bottom)** The IAV delivery platform enables a parallel process where wild-type cells are expanded in parallel to viral rescue and expansion. Alternatively, bioreactors can be maintained at production titers. Once at production titers, the IAV gene delivery reprograms the whole reactor for protein production.

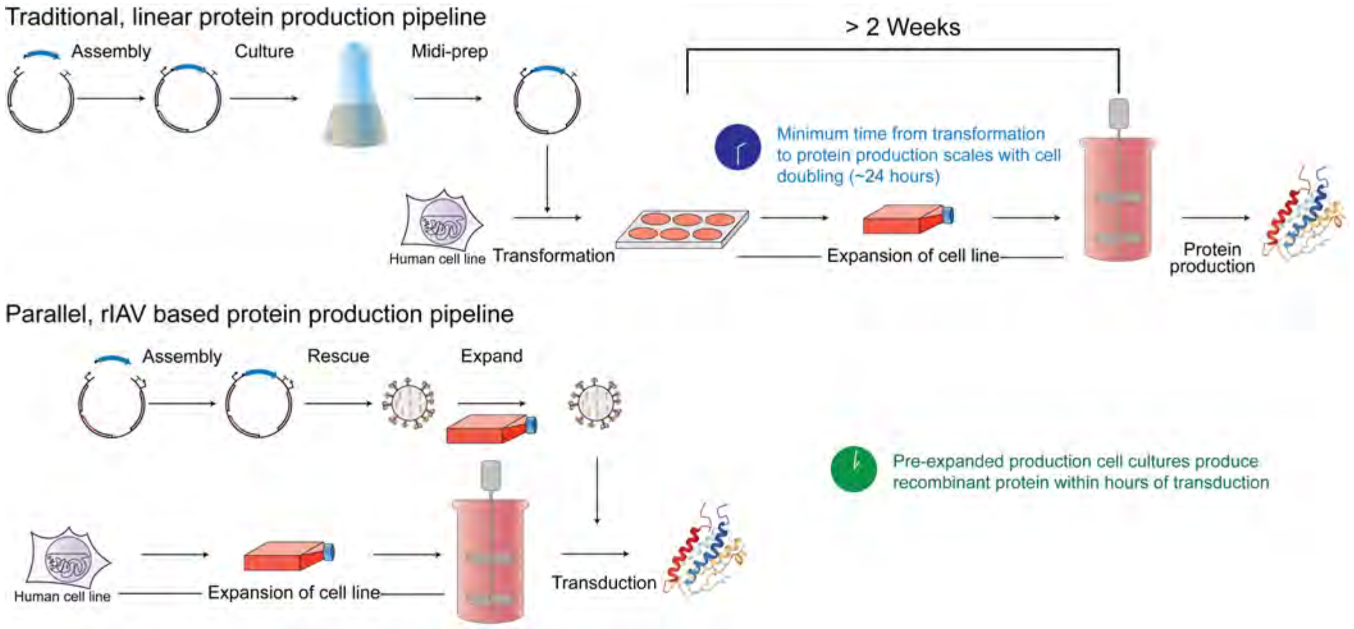


**Fig. 9**

DRAFT

### Supplementary Note 6: 5′ to 3′ vRNA sequence of SARS-CoV-2-S(Wuhan-Hu-1)-sciIAV

AGTAGAAACAAGGGTGTTTTTCCTCATATTTCTGAAATTCTAATCTCAGATGCATATTCTGCACTGCAAAGAT
CCATTAGAACACATCCAGAAACTGATTGCCCCCAGGGAGACCAAAAGCACCAGTGAACTGGCGACAGTTGAGT
AGATCTCGAGTTACACCGTTGCAGGCGCATTGAGGAGCTCAAAAGACAGGACTACAACCCGGTAGGGTTGATA
CCCGATTCCAGTGGTCGTATAGAACCCATAATCGTTAAGTGGCCAGTAGCAATTCAGAGCTGGGGGTGTGCAA
GGTTTTCCATCTGGACTAAAGGGAACATTTGATATGTCTCGCTCAAAAGGTCTCAATTTTCCATGTCTCAAAT
ACCTGTACTTGTAGTTATAGTTACCAGTAGATGTCGCATCAATGTTCCTGGTATTCCAAGCGAGAACACAACC
CATAAAATCATCAGGAAGTTTGTAGTTATAATCAGCGATCACTCCCGTTTGTCCTGGAGCGATCTGTCTAACG
TCGTCGCCTTTGACTACAAAGGAATCGGCGTATACATTGCTGAAGCAAAGATCATTCAGCTTAGTAGCGGAGA
CTCCGTAGCATTTGAAGGTAGAAAAGAAAGTAGAGTTATAAAGCACACTGTAATCTGCGACACAGTTGCTGAT
CTTCTTTCTTTCCCACGCATAAACGGAAGGGAACTTTGTGGCGTTAAAAACTTCGCCGAAAGGACAGAGATTT
GTAATGTTAGCCCAAACCATTGGCCAGAGAAGCAAAAGCAGCAGGAGCAGCCACCACAGCCGCCACCACATGG
CTAGCCTATACAAATTGTGTCTGCAACTGCAGCTGCAAGTGCACTTAACAGGACCAGTAGGTTTGCCTTCAAT
TTGGTTGTTTTTATTTTCCCCTGCTTTTGCT

DRAFT

### Supplementary Note 7: 5′ to 3′ vRNA sequence of SARS-CoV-1-S(Tor2)-sciIAV

AGTAGAAACAAGGGTGTTTTTCCTCATATTTCTGAAATTCTAATCTCAGATGCATATTCTGCACTGCAAAGAT
CCATTAGAACACATCCAGAAACTGATTGCCCCCAGGGAGACCAAAAGCACCAGTGAACTGGCGACAGTTGAGT
AGATCTCGAGTTATACTGTGGCGGGAGCGTGAAGAAGCTCGAATGAAAGGACTACGACTCTGTACGGTTGGTA
TCCGACCCCATTGGTTGGCTGGAATCCATAACTTTGAAGTGGGAAGTAGCAGTTAAACCCCTCTACACCGTTG
CAAGGCGTGGAGCCTGCCTGGTAAATTTCTGTGCTTATGTCTCTCTCAAAGGGTTTAAGATTTGACTTCCTGA
ACAATCGATACAGGTAATTATAATTTCCGCCGACTTTGCTGTCAAGATTATTGGAGTTCCATGCGATCACACA
CCCCGTGAAATCGTCCGGCAGTTTATAGTTATAATCGGCGATTTTACCTGTTTGACCTGGCGCTATTTGTCTG
ACCTCATCACCTCGGATAACGAAAGAATCGGCGTATACGTTGGTGAAACACAGATCATTCAACTTCGTTGGGG
ATACGCCGTAGCATTTGAACGTGCTAAAGGATGCACTATTATAGAGCACGGAATAATCTGCGACACAATTACT
GATCCTCTTGCGATTCCATGCATAAACACTTGCAAATCGTGTGGCGTTAAAAACTTCGCCGAAGGGACACAGG
TTAGTAATATTCGGGAACCTGACAATGCTTTCCGTCGGCTGAACCCGGAAATTGCTCGTTTGATATATACCTT
TTTCAACTGTGAAGGATTTCAAGGTACACTTAGTCTCACTCAACGGATCGAGAGCACAGTCTACGGCGTCGGT
TATTGTGCCATTCTCATTGTACTTCAGCAAGAACGTTCGCGGTTGAAGGTAACCTACATAATAAGCAGCTGCG
CCCGCAGTCCAGCCGCTGGAACTATCTCCGGGGGTAAGATAACTGCGATGCAGCGCGAGCAGCGTCTGGAACC
GCGTAATGTTGGCCCAGACCATAGGCCACAAAAGGAGGAGGAGCAGCAGGAGCCACCAAAGCCTCCACCACAT
GGCTAGCCTATACAAATTGTGTCTGCAACTGCAGCTGCAAGTGCACTTAACAGGACCAGTAGGTTTGCCTTCA
ATTTGGTTGTTTTTATTTTCCCCTGCTTTTGCT

DRAFT

### Supplementary Note 8: 5′ to 3′ vRNA sequence of SARS-CoV-2-S(BA.1)-sciIAV

AGTAGAAACAAGGGTGTTTTTCCTCATATTTCTGAAATTCTAATCTCAGATGCATATTCTGCACTGCAAAGAT
CCATTAGAACACATCCAGAAACTGATTGCCCCCAGGGAGACCAAAAGCACCAGTGAACTGGCGACAGTTGAGT
AGATCTCGAGtcaGAAGTTTACACACTTGTTCTTGACCAAGTTGGTAGACTTCTTTGGGCCGCATACTGTTGC
TGGCGCATGAAGCAGTTCGAAGCTCAAAACTACTACTCTGTAAGGCTGATGTCCGACACCATATGTCGGTCGG
AATGAATATGAGCGAAGCGGGAAATAGCAATTGAATCCTGCGACACCGTTGCACGGCTTGTTTCCGGCTTGAT
AAATCTCGGTGGAGATATCCCTCTCGAAAGGCTTCAAATTGCTCTTTCTGAACAGCCTGTAGAGATAATTGTA
ATTGCCAGATACCTTGCTATCCAGCTTGTTTGAGTTCCAAGCGATGACGCACCCGGTGAAATCGTCGGGCAAT
TTGTAGTTGTAGTCGGCGATGTTCCCCGTCTGGCCGGGTGCTATCTGGCGCACCTCATCACCTCTAATCACGA
AGGAATCAGCATACACGTTGGTGAAACAGAGATCGTTGAGCTTGGTCGGAGAAACTCCGTAGCATTTAAAGGT
AAAGAACGGTGCGAGGTTATAGAGGACAGAGTAGTCGGCAACGCAGTTACTAATTCTCTTTCTGTTCCACGCA
TACACGCTGGCAAACCGCGTTGCGTTAAAGACCTCATCAAATGGACACAAGTTTGTGATATTCGGGAACCGAA
CGATACTCTCTGTCGGTTGGACCCGGTGGTGGTGATGATGGTGGGCTTCGGCCACGGCAATACAGATCAGGGC
GAACAGgACCTTCACgCCCATGGCTAGCCTATACAAATTGTGTCTGCAACTGCAGCTGCAAGTGCACTTAACA
GGACCAGTAGGTTTGCCTTCAATTTGGTTGTTTTATTTCCCCTGCTTTTGCT

DRAFT

### Supplementary Note 9: 5′ to 3′ vRNA sequence of MBL-sciIAV

AGTAGAAACAAGGGTGTTTTTCCTCATATTTCTGAAATTCTAATCTCAGATGCATATTCTGCACTGCAAAGAT
CCATTAGAACACATCCAGAAACTGATTGCCCCCAGGGAGACCAAAAGCACCAGTGAACTGGCGACAGTTGAGT
AGATCTCGAGTTATATGGGGAATTCGCAGACAGCCAAATGGCTTGTACTGCAGGGCACATCGTTCCACTGACC
ATTCTTCAGCAGGAGCACGCAATCCTCGTCGGAACCCGCATTATTGGGCTCGCCTTCATTCCAGTTAGTGTAT
GTGAGTCGGTTGCCCGTGAGATCGACGAATTGCCCCTCTGTCTTTTCATCTGTTATCCCGAGAAAAGCCTCCT
CCTTTATGAGGTTTTGGATGGCGCCGTTTTCAGCGGCATTTCGCGGAGTTGCAACGCTGGCCTGGAATTTAAC
ACACAGGGCCTTAACCTTCTCAAACGTCATTATTTCACCATTGGTCAAAAAGAATTTGTTCCCTACCTGCTTG
CCCAGACTAAATGTCAGCCACTTTTTAATGCGTGCCATTTCGGTTTGCAAGGCTTTTCGCTCGCTCGCAGCCA
AAGAGGAGTCCCCGTCGGGAGATTTGCCGGGATCTCCTTTCTGACCCTTAGGACCCGGGCTCCCGCTCGGTCC
AGGATTACCAGGTGGTCCCAGCTTTCCTGGAGGACCTTGCAATCCCCTAAGGCCTTGACCTGGCTCTCCTTTC
TCCCCTTTGGTACCATCTCTGCCGTCCTTGCCCGGGAATCCATTGATACCAGGTGAACTGCACGCTATCACCG
CTGGACAAGTCTTCTGTGCATCTTCGCAAGTGACTGTTTCAGAATAAGATGCCGCTACCATGGAAAGCAACAA
CAGCGGCAGTGACGGGAAAAGGCTCATGGCTAGCCTATACAAATTGTGTCTGCAACTGCAGCTGCAAGTGCAC
TTAACAGGACCAGTAGGTTTGCCTTCAATTTGGTTGTTTTTATTTTCCCCTGCTTTTGCT

DRAFT

### Supplementary Note 10: Full Western Blots

Representative uncropped Western Blots from Figure 2f. a) Western Blot of protein in supernatant stained by anti-His, anti-SARS-CoV, and anti-MBL antibodies. b) Western Blot of protein in cell lysate stained by anit-His, anti-SARS-CoV, and anti-MBL antibodies.

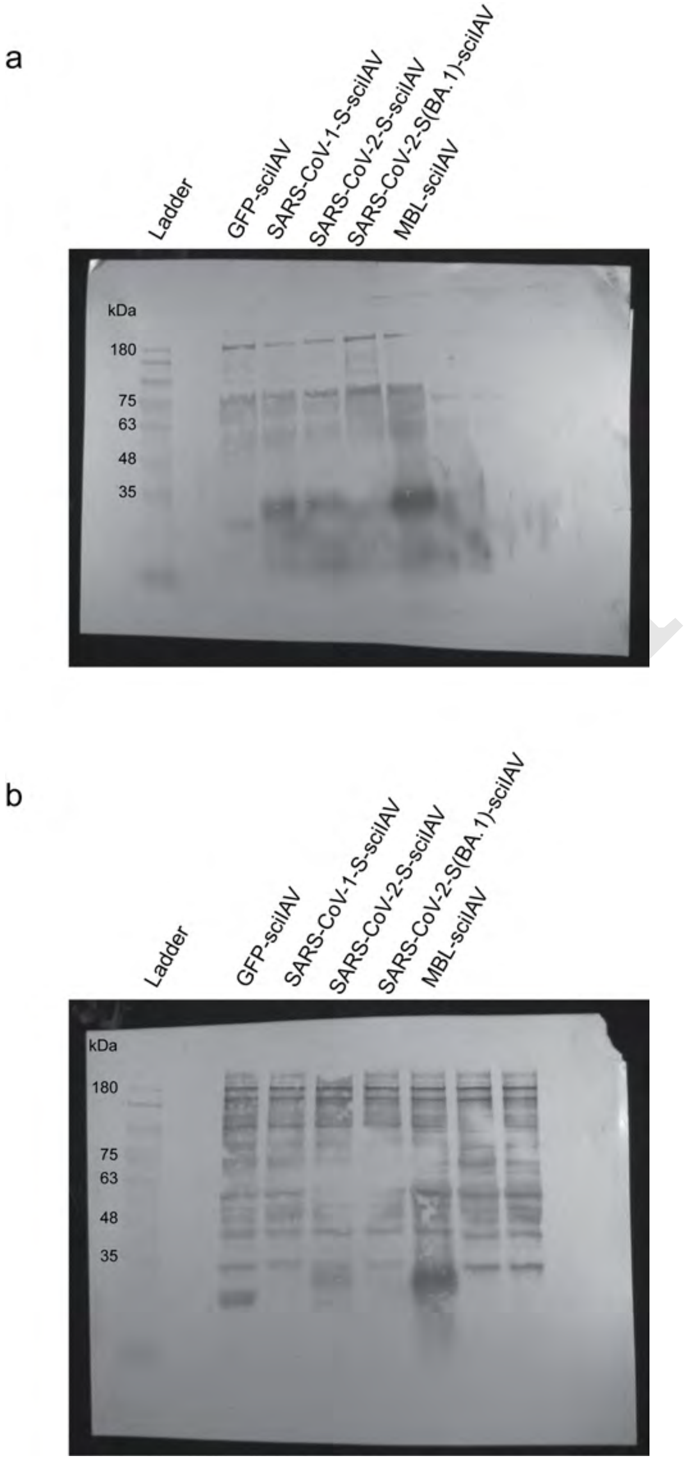


**Fig. 10**

### Supplementary Note 11: Entropy and Mutual Information for plasmids in library 1

a) Amino acid entropy in the modified region of library 1 plasmids b) Nucleotide entropy in the modified region of library 1 plasmids c) Nucleotide pairwise mutual information in modified region of library 1 plasmids

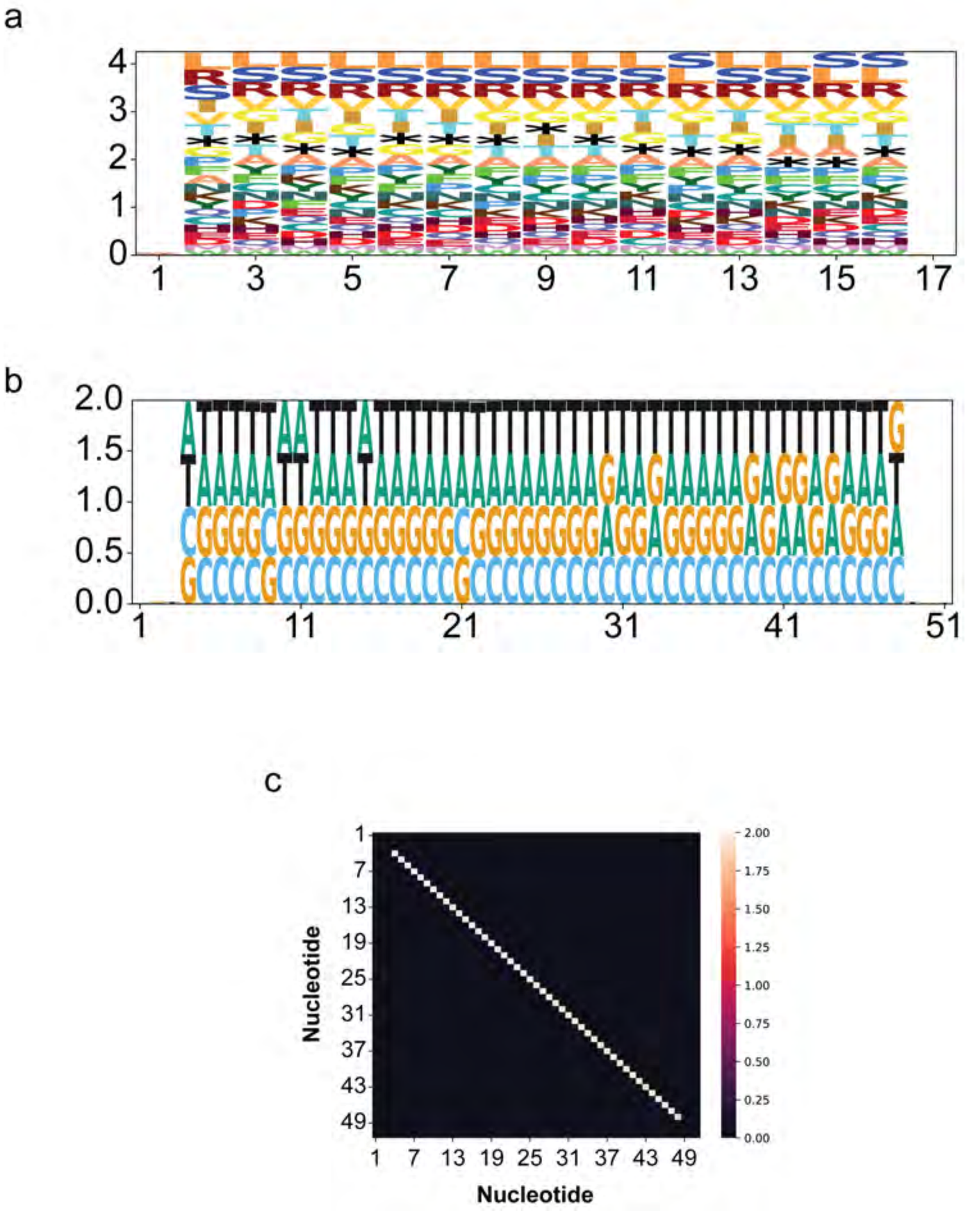


Fig. 11

### Supplementary Note 12: Jensen-Shannon Divergence at nucleotide level

JSD was calculated for each base at each nucleotide. a) Plasmid Library vs. every other library. b) Packaging Library vs. every other library. c) Propagation Library vs. every other library. d) Sorted Library vs. every other library. e) High Expression Library vs. every other library. f) Medium Expression Library vs. every other library. g) Low Expression Library vs. every other library. h) GFP Negative Library vs. every other library.

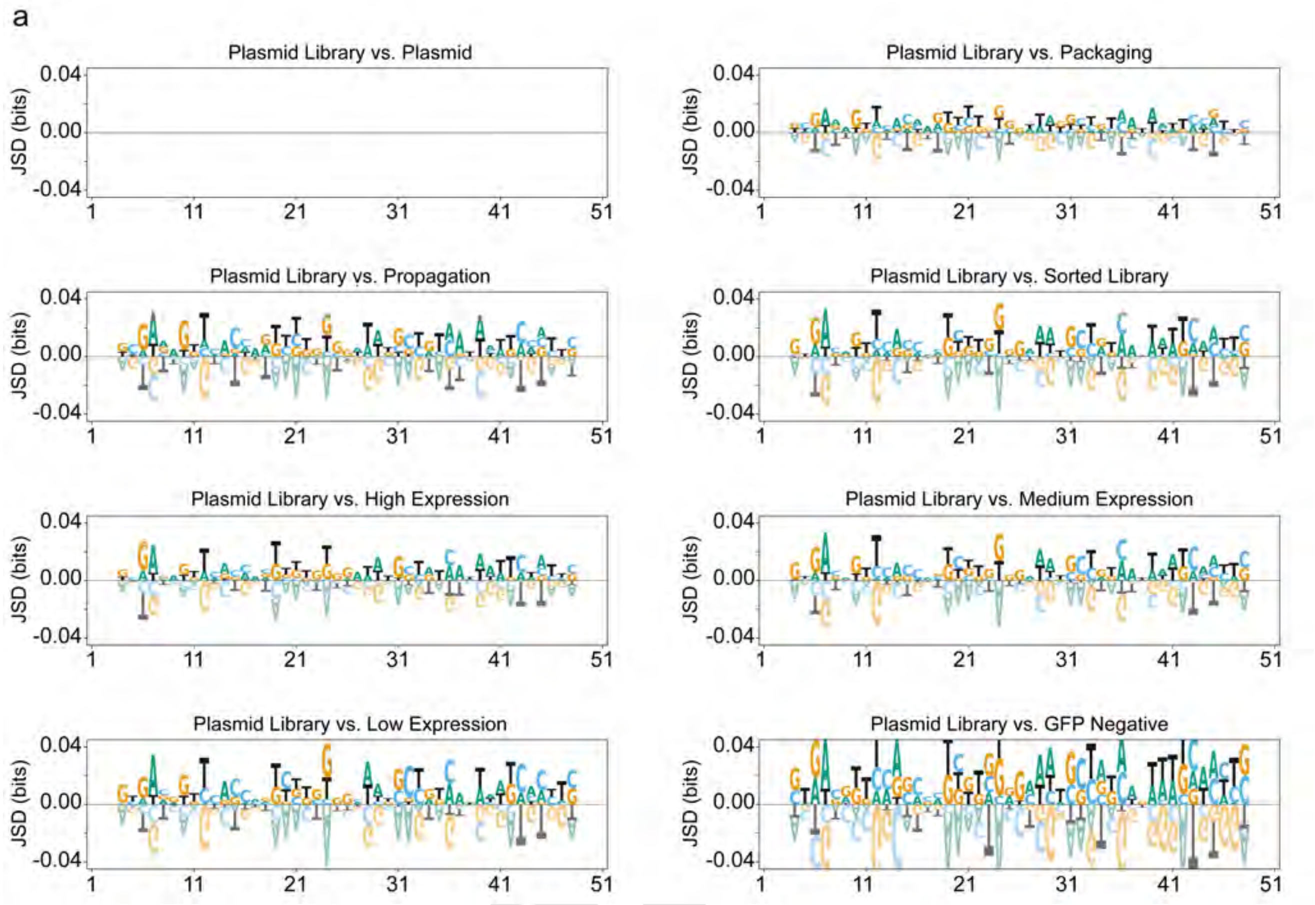


**Fig. 12**

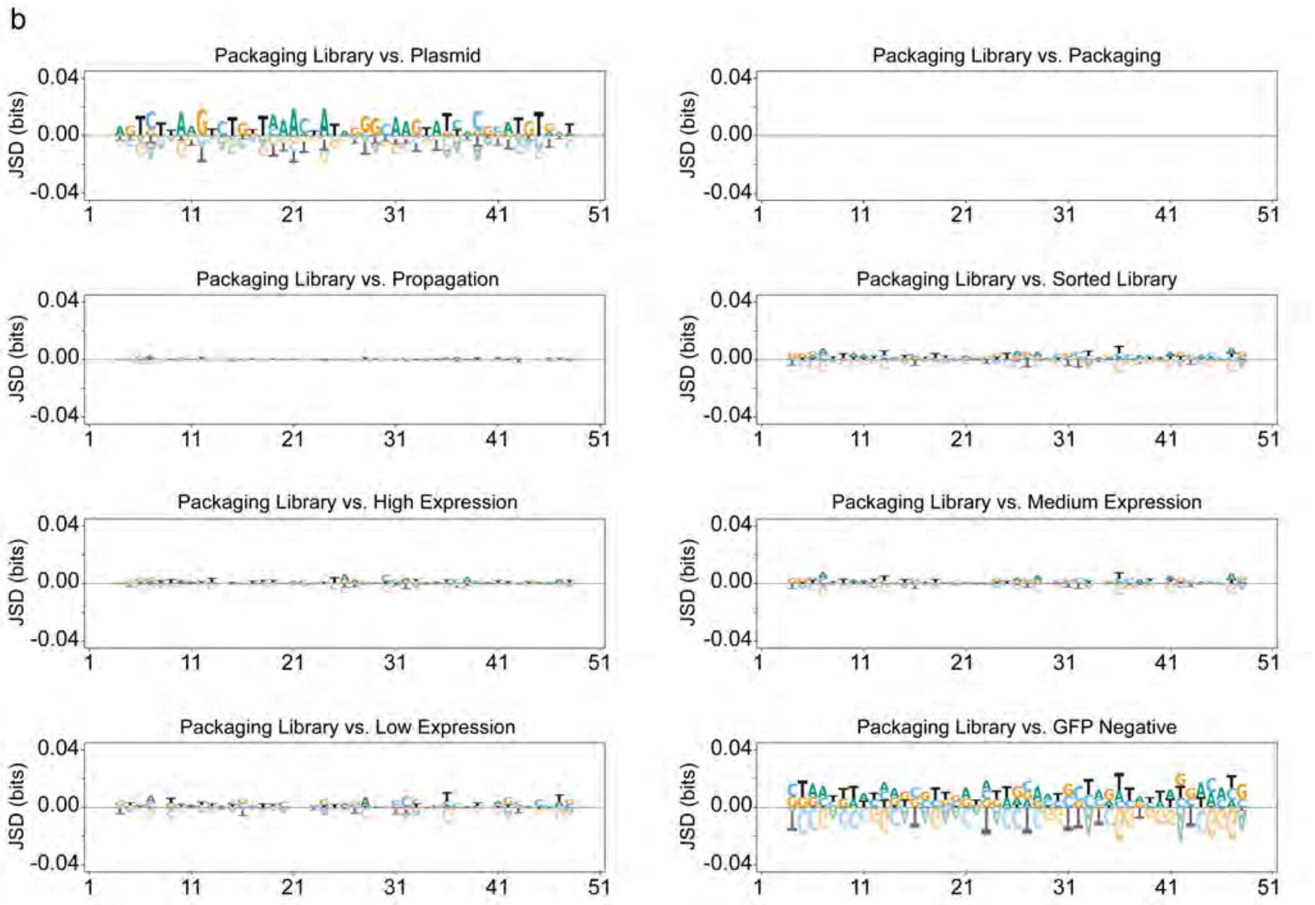
b
Packaging Library vs. Plasmid
Packaging Library vs. Packaging
Packaging Library vs. Propagation
Packaging Library vs. Sorted Library
Packaging Library vs. High Expression
Packaging Library vs. Medium Expression
Packaging Library vs. Low Expression
Packaging Library vs. GFP Negative
JSD (bits)
0.04
0.00
-0.04
1
11
21
31
41
51

**Fig. 13**

DRAFT

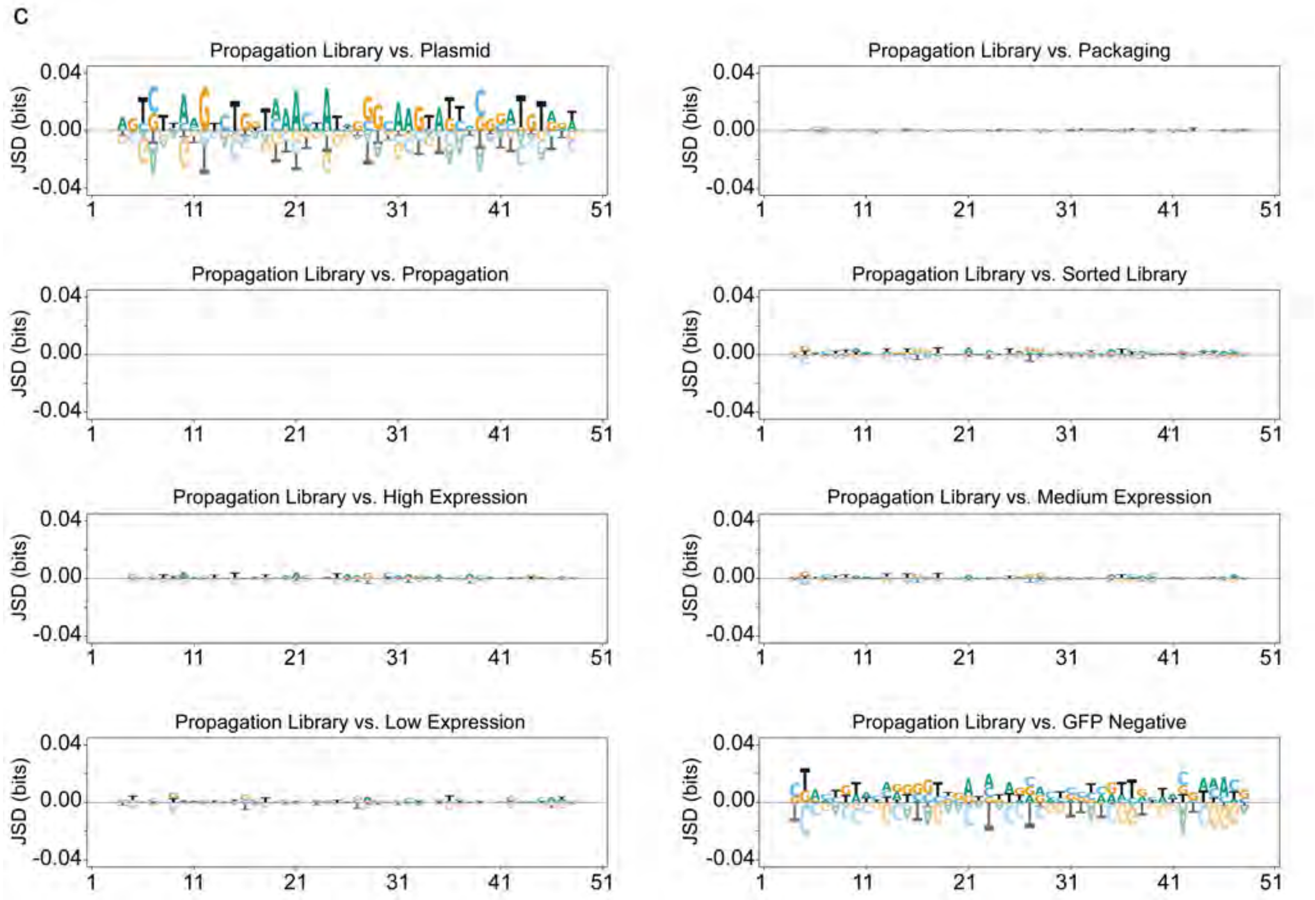
C
Propagation Library vs. Plasmid
Propagation Library vs. Packaging
Propagation Library vs. Propagation
Propagation Library vs. Sorted Library
Propagation Library vs. High Expression
Propagation Library vs. Medium Expression
Propagation Library vs. Low Expression
Propagation Library vs. GFP Negative
JSD (bits)
0.04
0.00
-0.04
1
11
21
31
41
51

**Fig. 14**

DRAFT

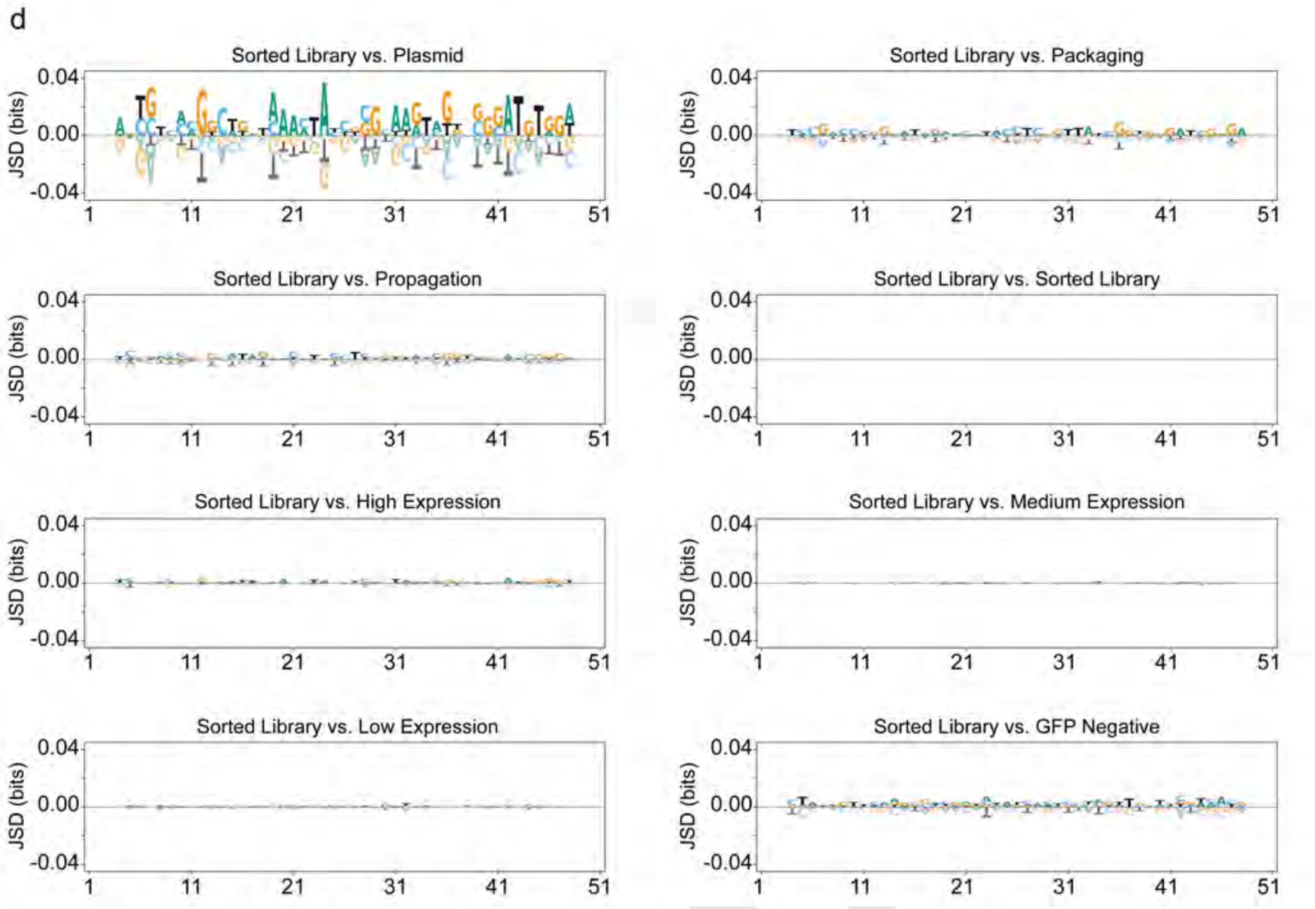
d
Sorted Library vs. Plasmid
Sorted Library vs. Packaging
Sorted Library vs. Propagation
Sorted Library vs. Sorted Library
Sorted Library vs. High Expression
Sorted Library vs. Medium Expression
Sorted Library vs. Low Expression
Sorted Library vs. GFP Negative
JSD (bits)
0.04
0.00
-0.04
1
11
21
31
41
51

**Fig. 15**

DRAFT

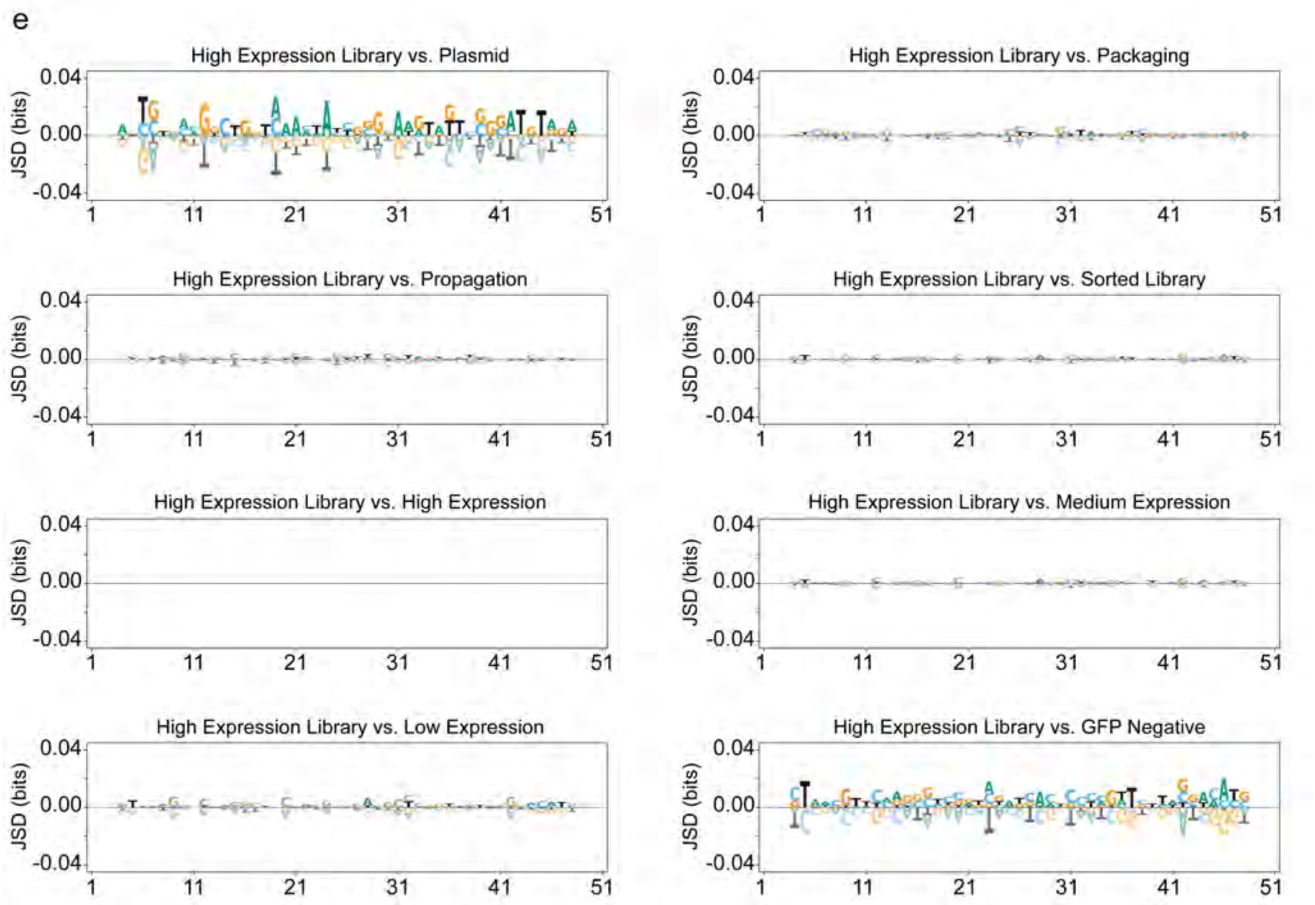
e
High Expression Library vs. Plasmid
High Expression Library vs. Packaging
High Expression Library vs. Propagation
High Expression Library vs. Sorted Library
High Expression Library vs. High Expression
High Expression Library vs. Medium Expression
High Expression Library vs. Low Expression
High Expression Library vs. GFP Negative
JSD (bits)
0.04
0.00
-0.04
1
11
21
31
41
51

**Fig. 16**

DRAFT

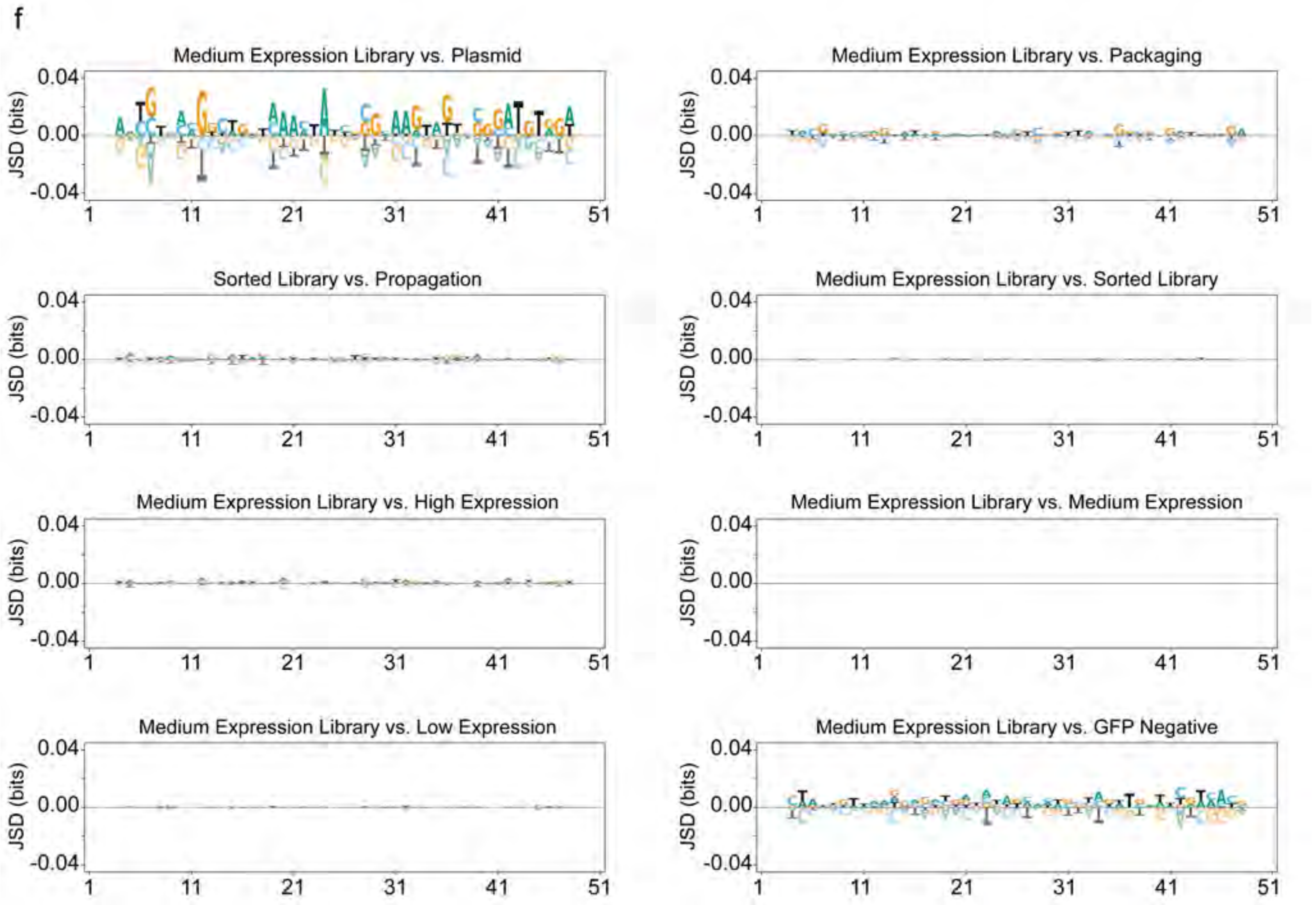
f
Medium Expression Library vs. Plasmid
Medium Expression Library vs. Packaging
Sorted Library vs. Propagation
Medium Expression Library vs. Sorted Library
Medium Expression Library vs. High Expression
Medium Expression Library vs. Medium Expression
Medium Expression Library vs. Low Expression
Medium Expression Library vs. GFP Negative
JSD (bits)
0.04
0.00
-0.04
1
11
21
31
41
51

**Fig. 17**

DRAFT

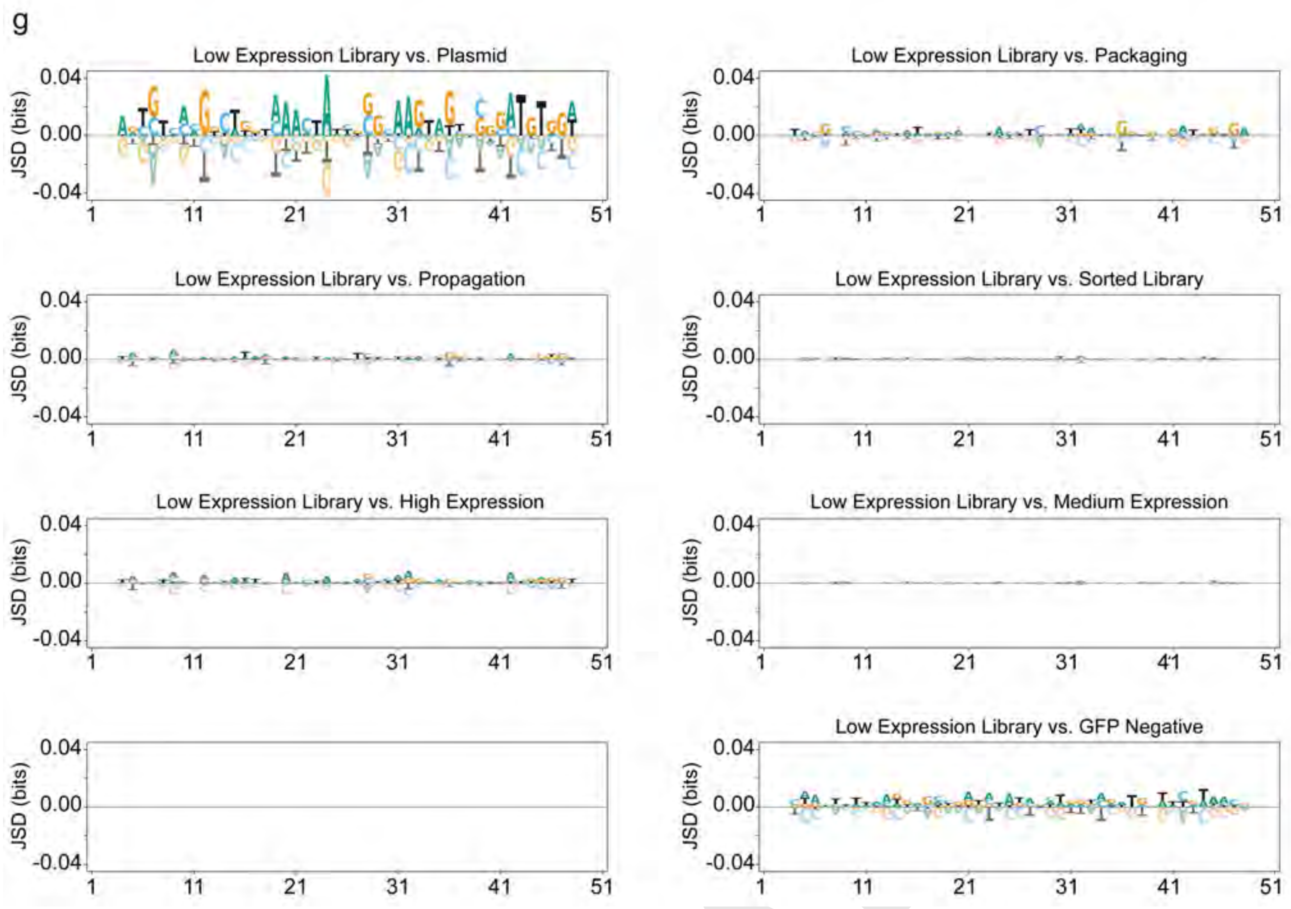
g
Low Expression Library vs. Plasmid
Low Expression Library vs. Packaging
Low Expression Library vs. Propagation
Low Expression Library vs. Sorted Library
Low Expression Library vs. High Expression
Low Expression Library vs. Medium Expression
Low Expression Library vs. GFP Negative
JSD (bits)
0.04
0.00
-0.04
1
11
21
31
41
51

**Fig. 18**

DRAFT

h

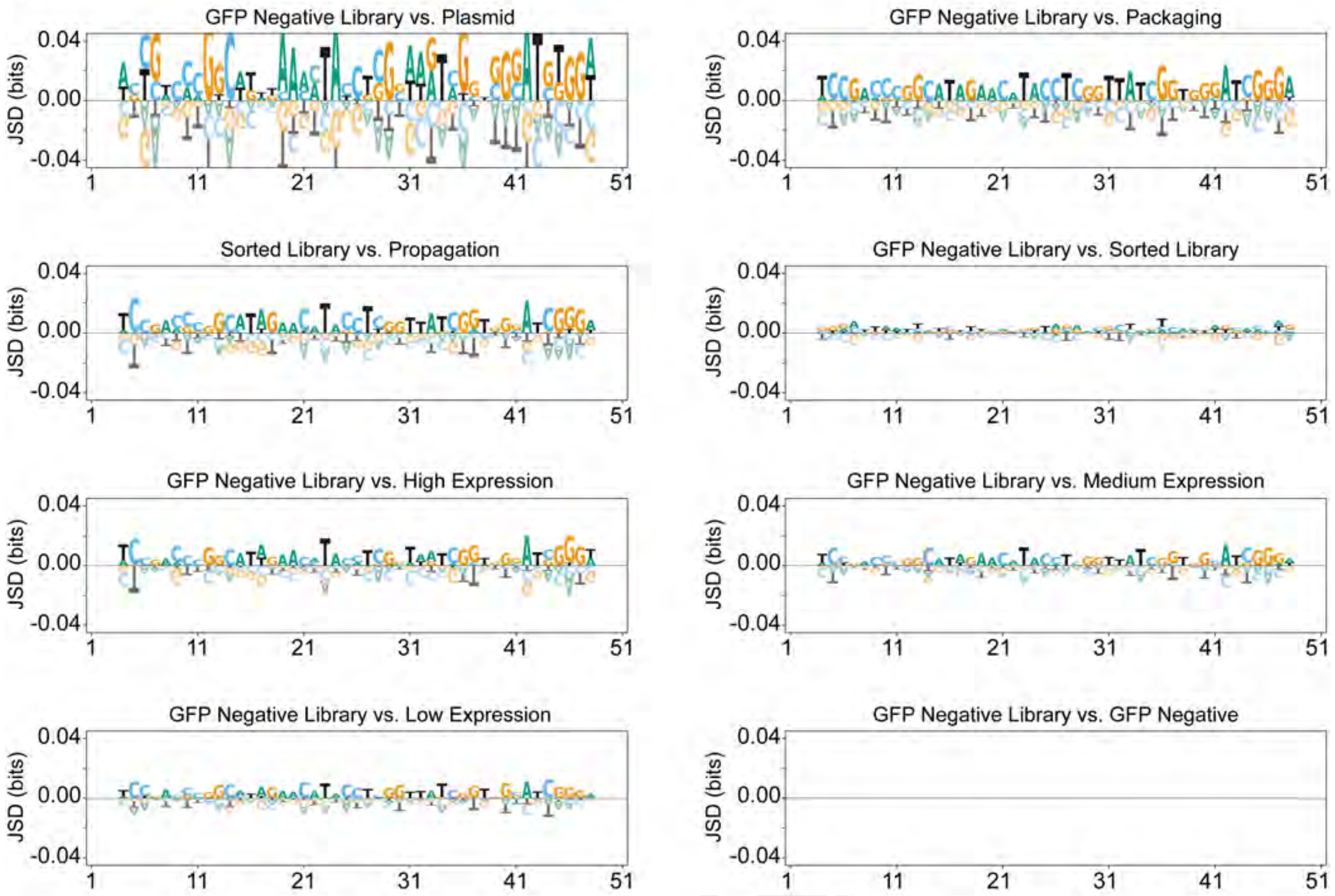


**Fig. 19**

DRAFT

### Supplementary Note 13: Entropy and Mutual Information for packaging library

a) Amino acid entropy in the modified region of library 1 b) Nucleotide entropy in the modified region of library 1 c) Nucleotide pairwise mutual information in modified region of library 1.

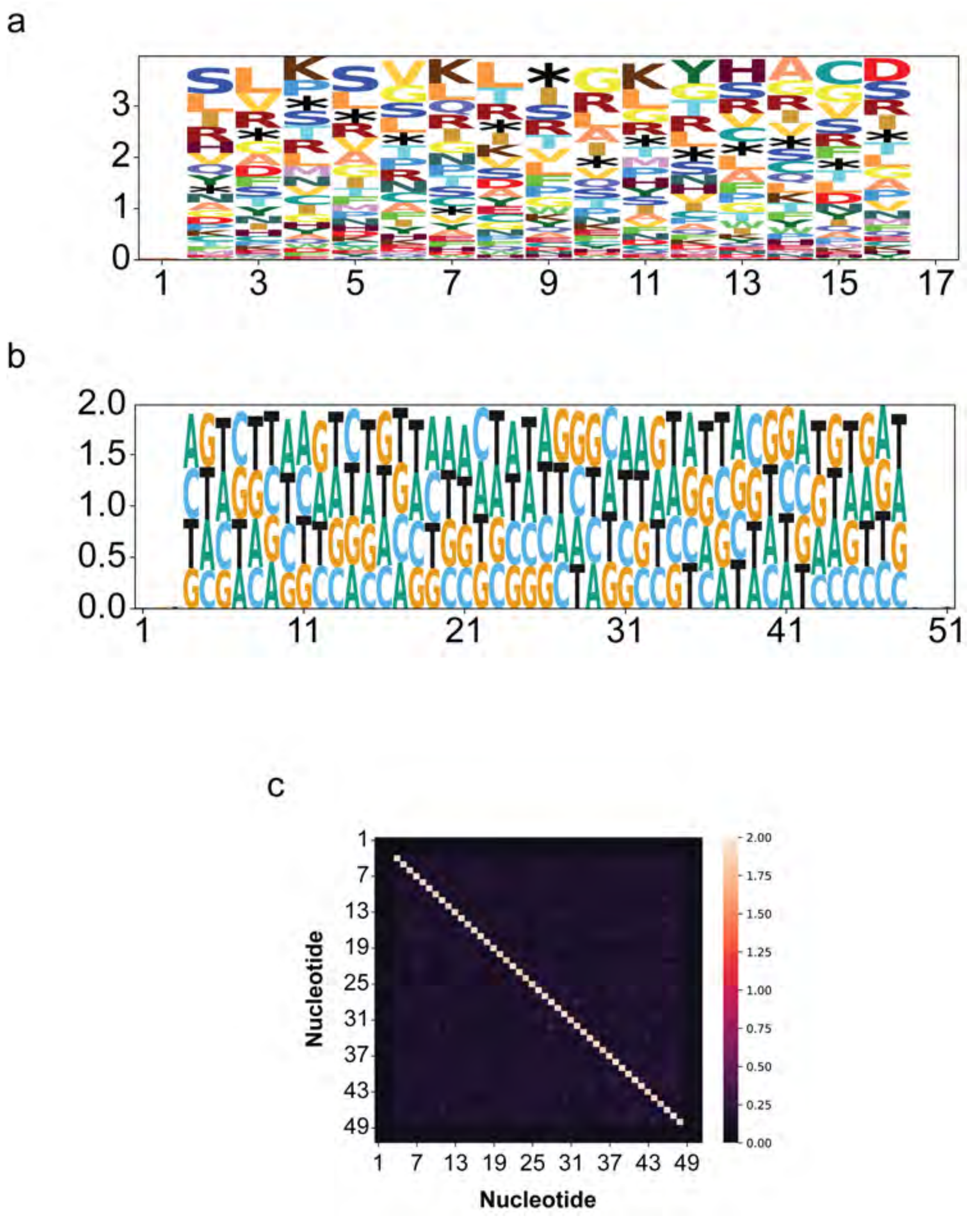


**Fig. 20**

### Supplementary Note 14: Entropy and Mutual Information for propagation library

a) Amino acid entropy in the modified region of library 1 b) Nucleotide entropy in the modified region of library 1 c) Nucleotide pairwise mutual information in modified region of library 1.

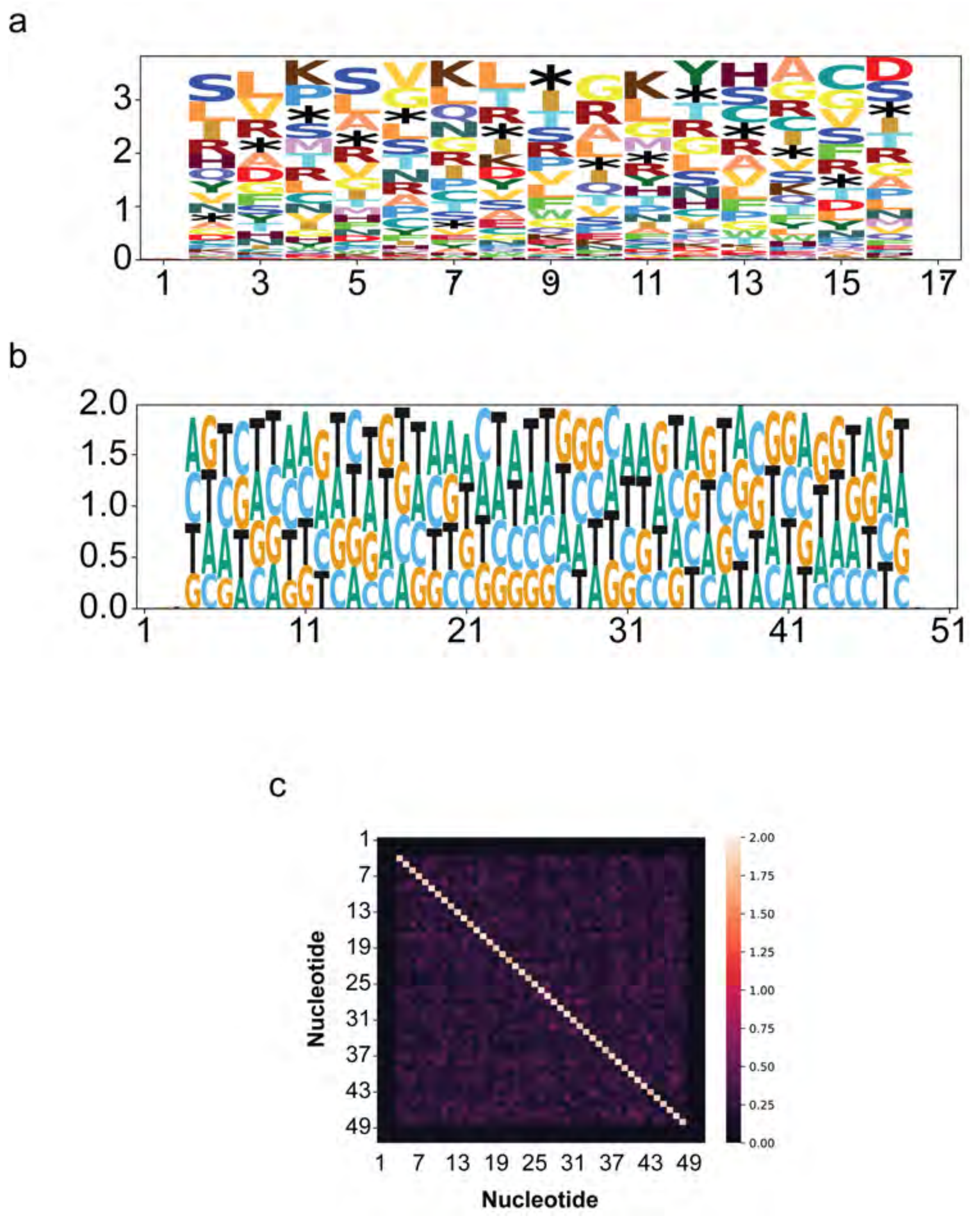


**Fig. 21**

### Supplementary Note 15: Entropy and Mutual Information for sorted library

a) Amino acid entropy in the modified region of library 1 b) Nucleotide entropy in the modified region of library 1 c) Nucleotide pairwise mutual information in modified region of library 1.

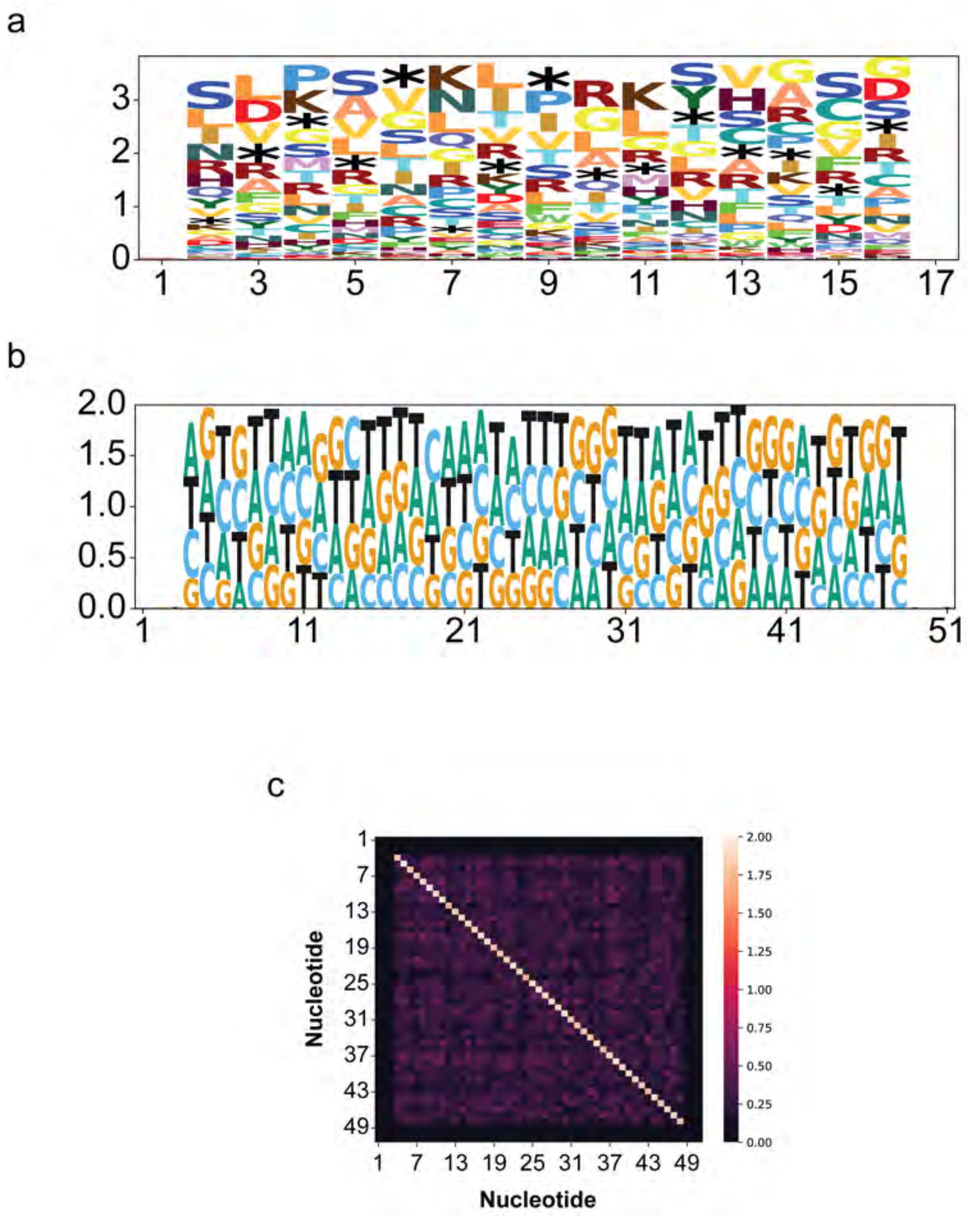


Fig. 22

### Supplementary Note 16: Entropy and Mutual Information for high expression library

a) Amino acid entropy in the modified region of library 1 b) Nucleotide entropy in the modified region of library 1 c) Nucleotide pairwise mutual information in modified region of library 1.

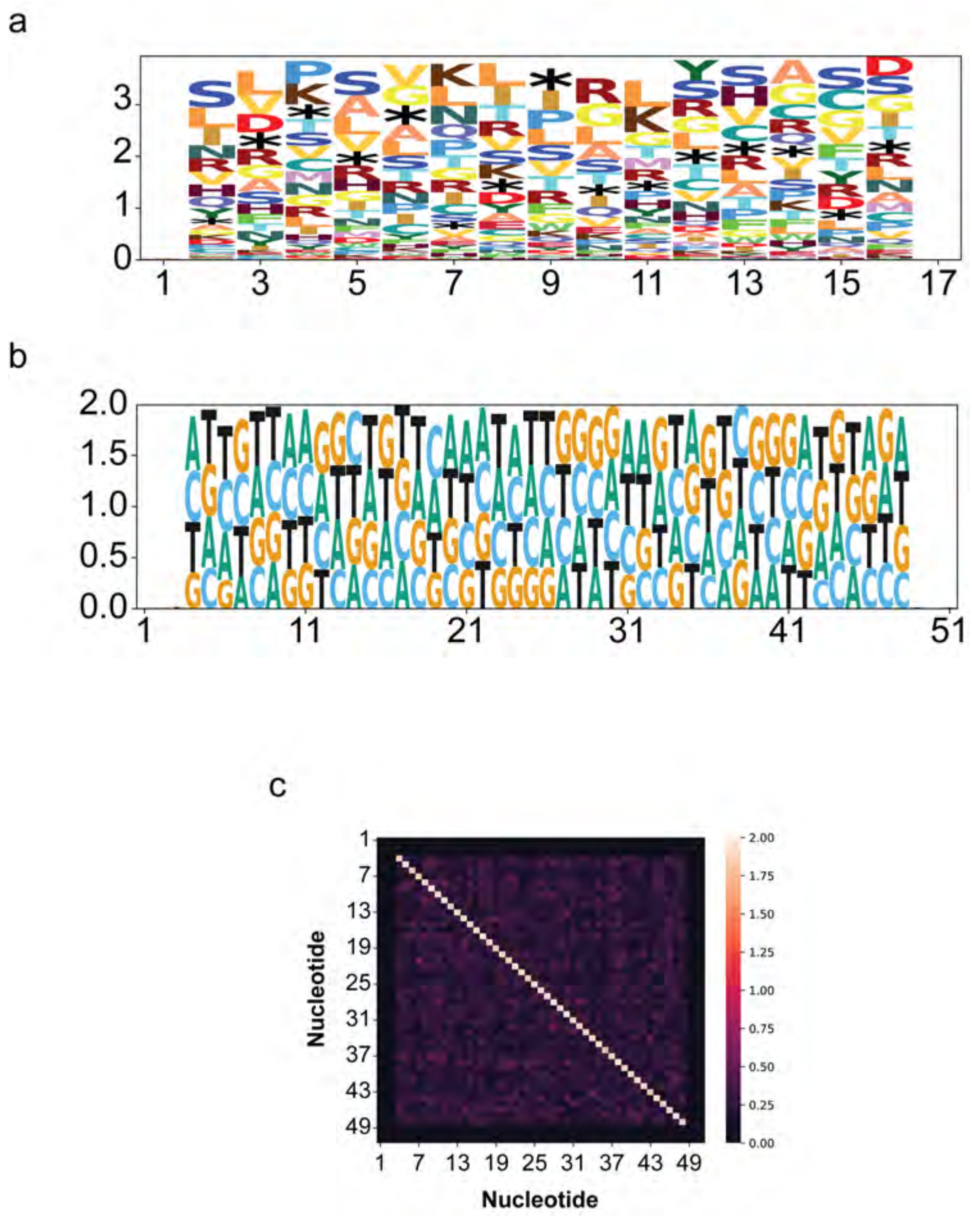


**Fig. 23**

### Supplementary Note 17: Entropy and Mutual Information for medium expression library

a) Amino acid entropy in the modified region of library 1 b) Nucleotide entropy in the modified region of library 1 c) Nucleotide pairwise mutual information in modified region of library 1.

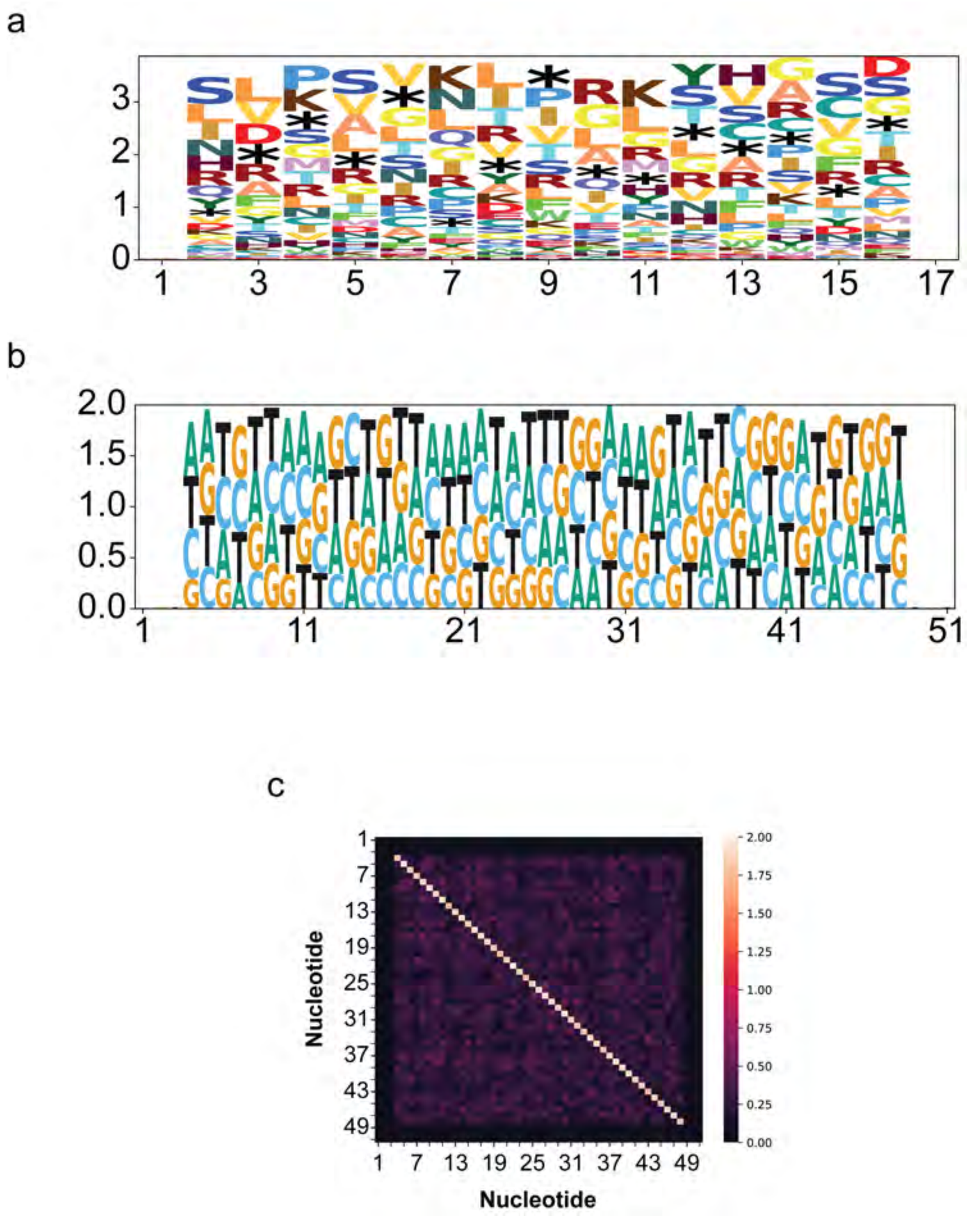


Fig. 24

### Supplementary Note 18: Entropy and Mutual Information for low expression library

a) Amino acid entropy in the modified region of library 1 b) Nucleotide entropy in the modified region of library 1 c) Nucleotide pairwise mutual information in modified region of library 1.

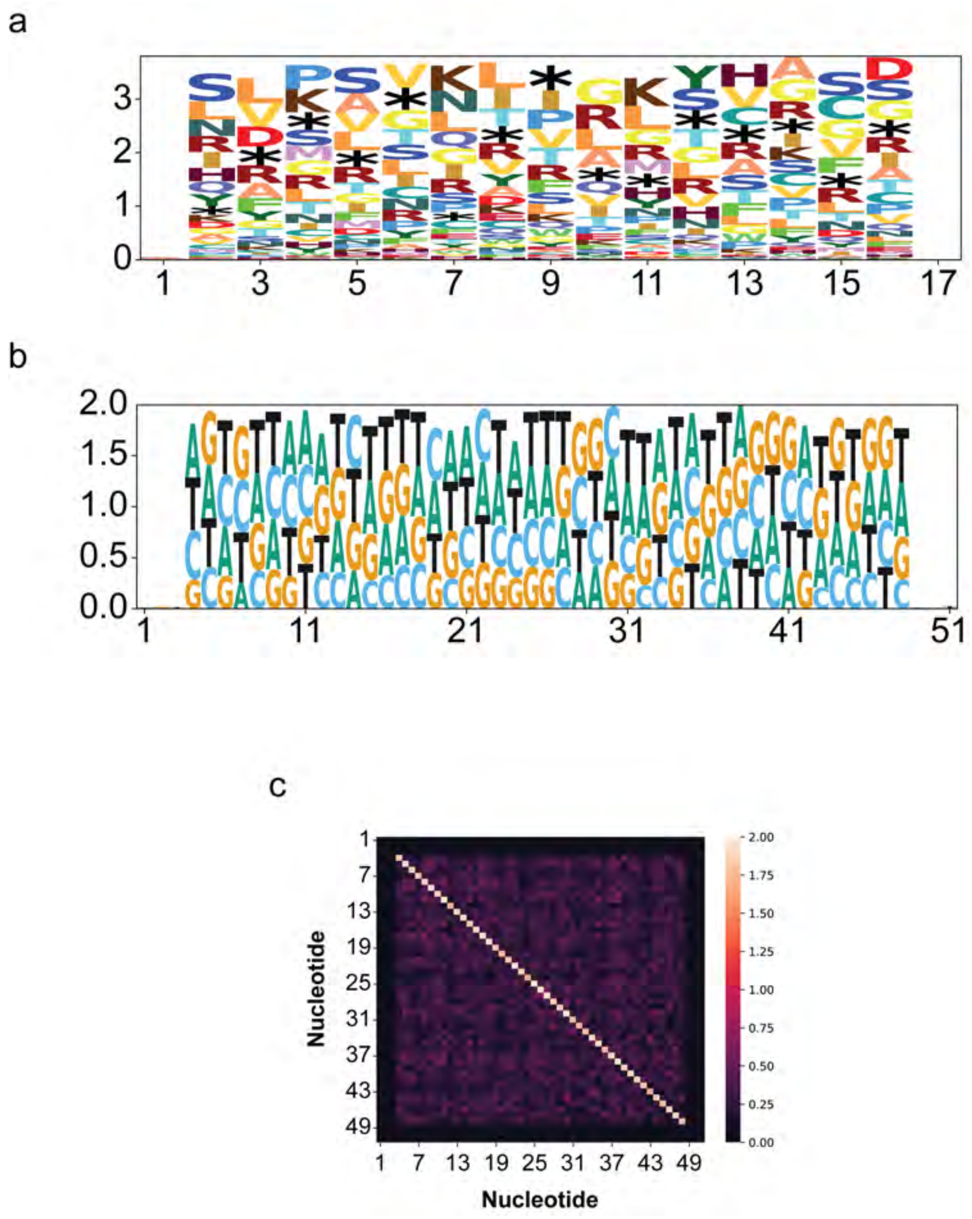


**Fig. 25**

### Supplementary Note 19: Entropy and Mutual Information for GFP negative library

a) Amino acid entropy in the modified region of library 1 b) Nucleotide entropy in the modified region of library 1 c) Nucleotide pairwise mutual information in modified region of library 1.

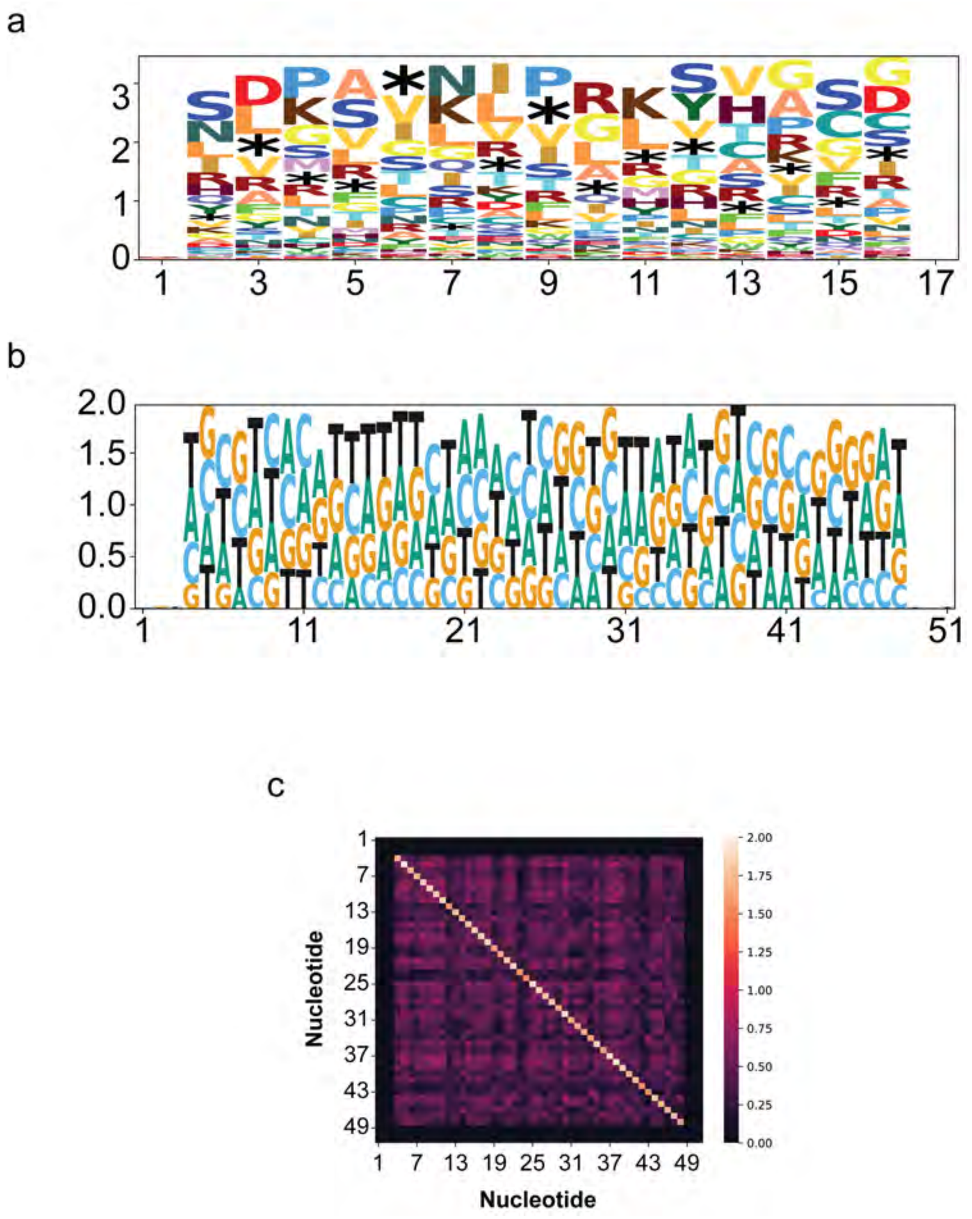


**Fig. 26**

| Primer Name | Sequence (5′ - 3′) | Target RNA | Strand Reverse Transcribed |
|---|---|---|---|
| oAS557 | GCTCAGGTGCATGTACCACT | Lysostaphin | (+) |
| oAS558 | AATTACGGCGGAGGCAATCA | Lysostaphin | (-) |
| oAS661 | CTACCCCGACCACATGAAGC | GFP | (-) |
| oAS662 | AAGAAGATGGTGCGCTCCTG | GFP | (+) |
| oAS378 | GGTGGTCTCCTCTGACTTCAACA | GAPDH (17) | (-) |
| oAS379 | GTTGCTGTAGCCAAATTCGTTGT | GAPDH (17) | (+) |

928 # Supplementary Note 20: Primers used for qRT-PCR in Figure 1d

DRAFT

| Primer Name | Sequence (5′ - 3′) | Target RNA | Strand Reverse Transcribed |
|---|---|---|---|
| oAS344 | AGTAGAAACAAGGGTGTTTTTCCTCATATTTCTGAA | HA Fragment | (+) |
| oAS345 | AGCAAAAGCAGGGGAAAATAAAAACAACCA | HA Fragment | (-) |

### Supplementary Note 21: Primers used for generating full length HA fragment cDNA

DRAFT

| Library | Primer Name<br>Reverse Primer Name | Forward Sequence (5′ - 3′)<br>Reverse Sequence (5′ - 3′) |
|---|---|---|
| Library 1: N-terminus | oAS719 | GACCTCCGAAGTTGGGGGGGAGCAAAAGCAGGNNNNNNNNNNNNNNNNNNNNGTGAGCAAGGGCGAGGAGC |
| Library 1: N-terminus | AS_rIAV_HA_F | GGCTAGCCTATACAAATTGTGTCTGC |
| Library 2: Comprehensive 3′UTR | oAS1200 | GACCTCCGAAGTTGGGGGGGAGCAAAAGCAGGNNNNNNNNNNNNNNNNNNNNATGGTGAGCAAGGGCGAGGAGC |
| Library 2: Comprehensive 3′UTR | AS_pDZ_R | CCCCCCCAACTTCGGAGG |
| Library 3: Restricted 3′UTR | oAS1200 | GACCTCCGAAGTTGGGGGGGAGCAAAAGCAGGNNNNNNNNNNNNNNNNNNNNATGGTGAGCAAGGGCGAGGAGC |
| Library 3: Restricted 3′UTR | AS_pDZ_R | CCCCCCCAACTTCGGAGG |

### Supplementary Note 22: Primers used for generating the viral libraries

DRAFT

| Primer Name | Sequence (5′ - 3′) |
|---|---|
| Forward Primer | ACACTCTTTCCCTACACGACGCTCTTCCGATCTctgacgacGCTCCTCGCCCTTGCTCAC |
| Reverse Primer | GACTGGAGTTCAGACGTGTGCTCTTCCGATCTctgacgacAGCAAAAGCAGG |

**Table 1.** Primers used for amplicon sequencing of viral libraries

### Supplementary Note 23: Primers used for sequencing the viral libraries

DRAFT

### Supplementary Note 24: Predicted vRNA, cRNA, and mRNA structures of the HA Segment

Schematics of predicted HA segment secondary structures using RNAfold (37). Bottom: full structure. Top: zoomed in view of inset. Red letters indicate the 3′ UTR. a) vRNA b) cRNA c) mRNA.

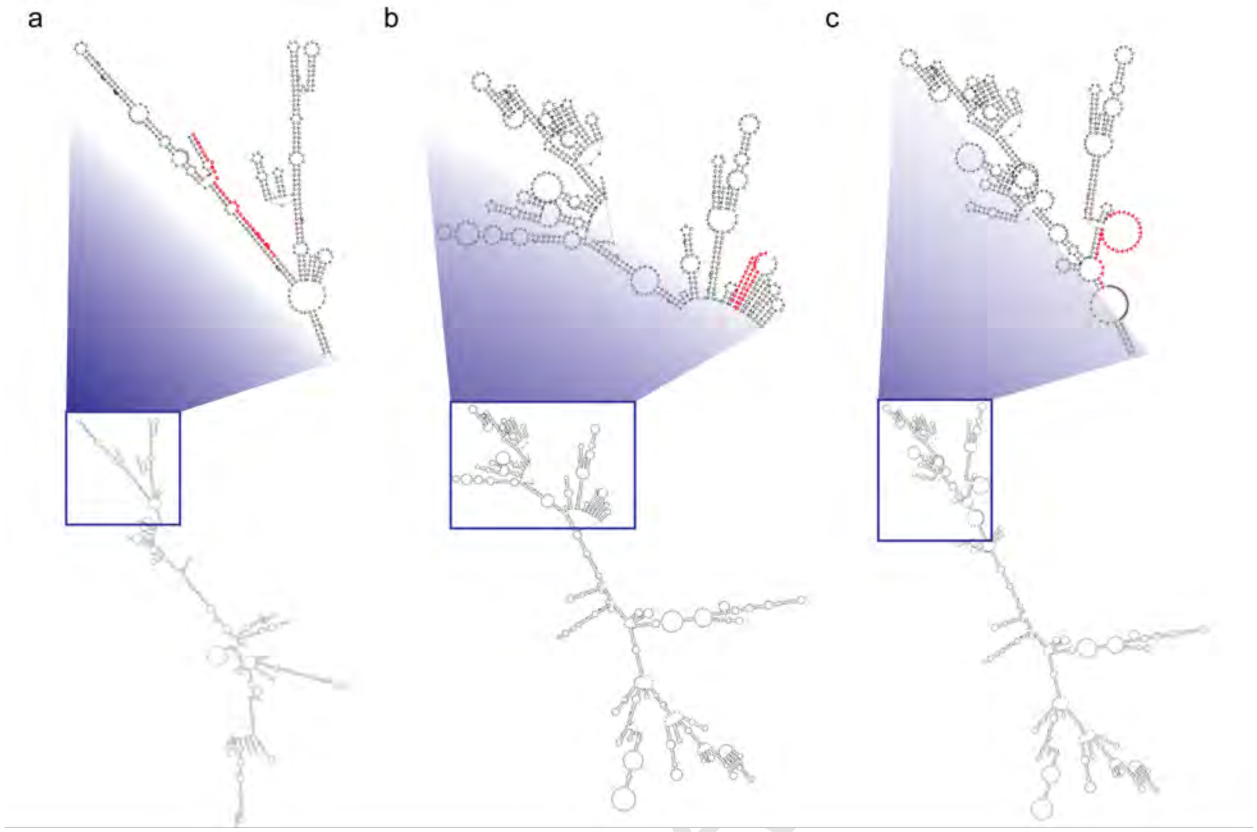


**Fig. 27**

DRAFT

### Supplementary Note 25: Zipf-Mandelbrot distribution of $3'$ UTR Library

Distribution of number of reads in the $3'$ UTR Library fit to a Zipf-Mandelbrot distribution using a Gauss-Newton algorithm. The distribution was found to be not statistically differnt from the data by the Kolmogorov–Smirnov test.

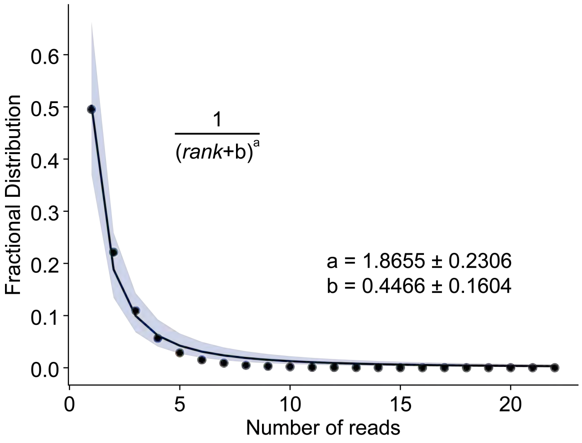


**Fig. 28**

DRAFT
